## Supplemental Figures and Tables for "Engineered reporter phages for rapid detection of *Escherichia coli, Klebsiella* spp., and *Enterococcus* spp. in urine"

### Supplementary Material – Meile et al. (2022)

#### Supplementary Figures

**Figure S1. Species distribution analysis of the Zurich Uropathogen Collection, related to Fig. 1.**

**Figure S2. An *Enterococcus*-adapted pSelect CRISPR-Cas9 system programmed to restrict wild-type phage plaque formation while allowing propagation of recombinant phages, related to Fig. 2.**

**Figure S3. Reporter phage specificity analysis, activity in blood, and clonal diversity within the Zurich Uropathogen Collection, related to Fig. 4.**

**Figure S4. Species distribution analysis of strains from the field evaluation, related to Fig. 5.**

**Figure S5. Definition of test parameters for a reporter phage-based companion diagnostic, related to Fig. 6.**

#### Supplementary Tables

**Table S1. The Zurich Uropathogen Collection.**

**Table S2. Strains used in this study.**

**Table S3. Bacteria isolated during the field evaluation from patient urine.**

**Table S4: Plasmids used and constructed during this study.**

**Table S5: Primer and synthetic DNA string sequences used in this study.**

### Supplementary Figures

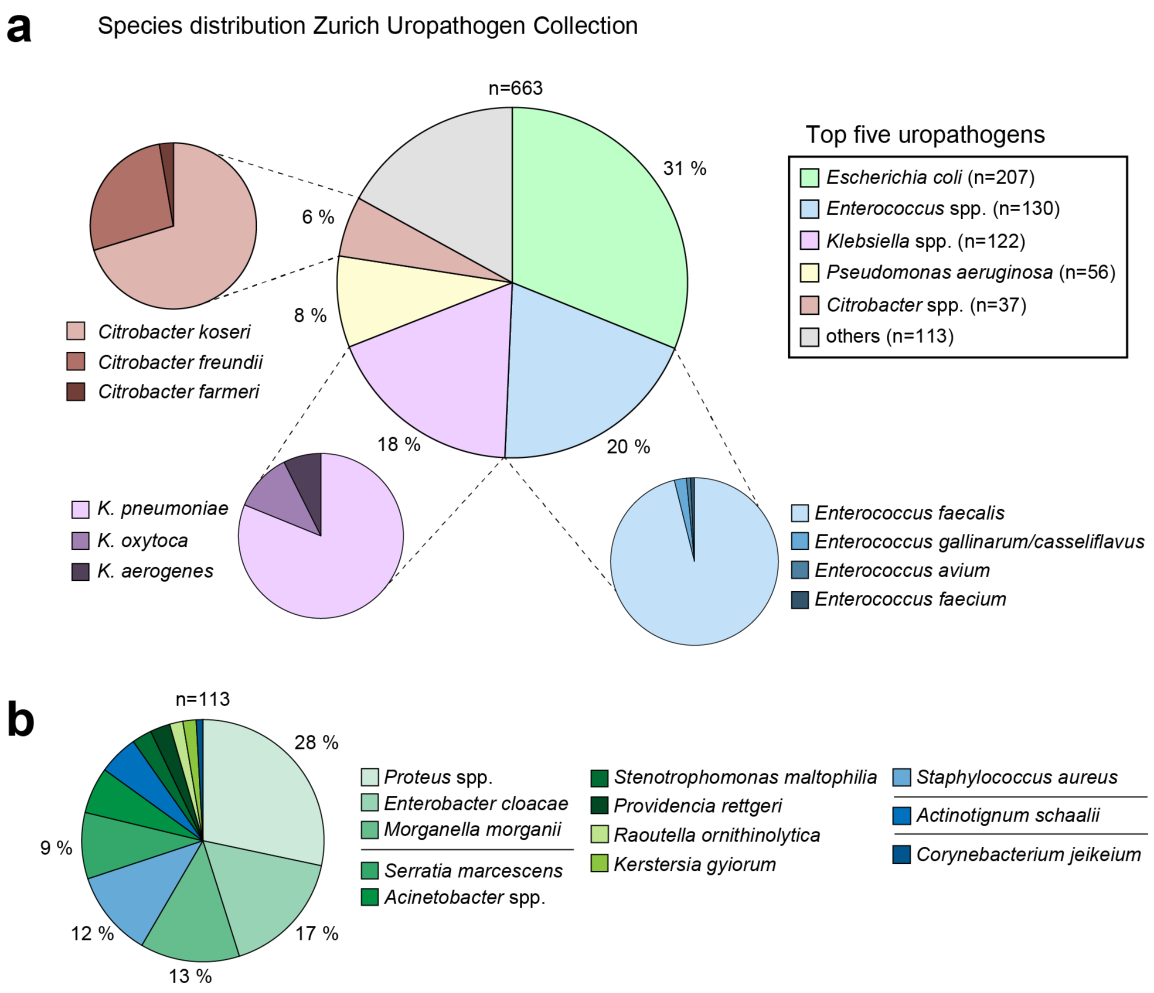


**Figure S1. Species distribution analysis of the Zurich Uropathogen Collection, related to Fig. 1.** (**a**) A total of 663 bacterial stains within the collection were analyzed with respect to their genus- and species distribution. Isolates that do not belong to the top five most prevalent uropathogens are shown in (**b**). *E. coli*, *Enterococcus* spp., and *Klebsiella* spp. isolates constitute the top three most prevalent uropathogens, collectively adding up to 69 % of isolates within the collection.

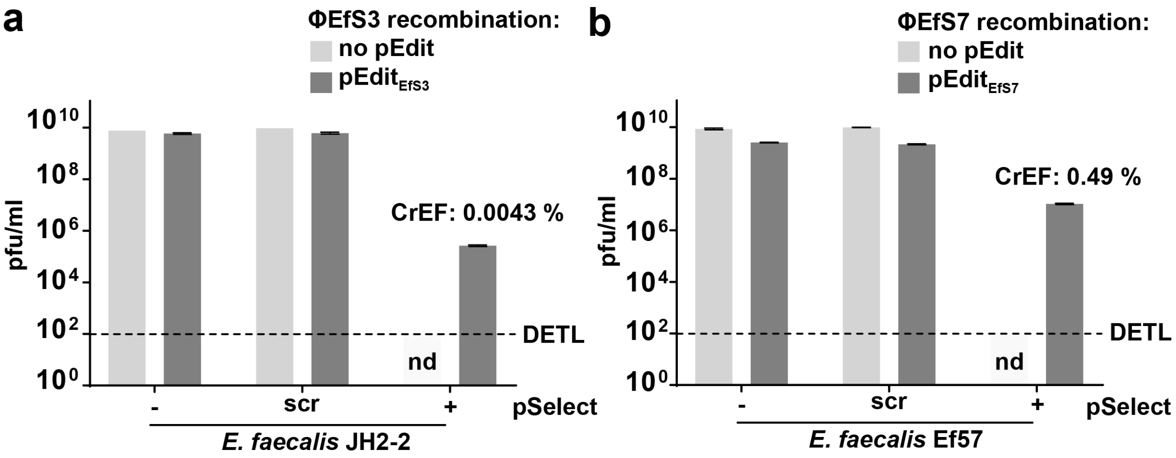


**Figure S2. An *Enterococcus*-adapted pSelect CRISPR-Cas9 system programmed to restrict wild-type phage plaque formation while allowing propagation of recombinant phages, related to Fig. 2.** Phage lysates were generated on hosts without (light grey) or with (dark grey) editing plasmid (homologous recombination). Plaquing efficiency of (**a**) EfS3 lysates and (**b**) EfS7 lysates on hosts with (+) or without (-, scr) CRISPR targeting through the pSelect system. Data is mean +/- SEM of biological triplicates. As wildtype phages are fully restricted in the presence of pSelect, the CRISPR-escape frequency ($PFU/mL_{+}$ / $PFU/mL_{scr}$) can be used as an approximation for the frequency of homologous recombination. scr = non-targeting crRNA; - = no pSelect; CrEF: CRISPR-escape frequency; DETL = detection limit for plaque formation; nd = not detected.

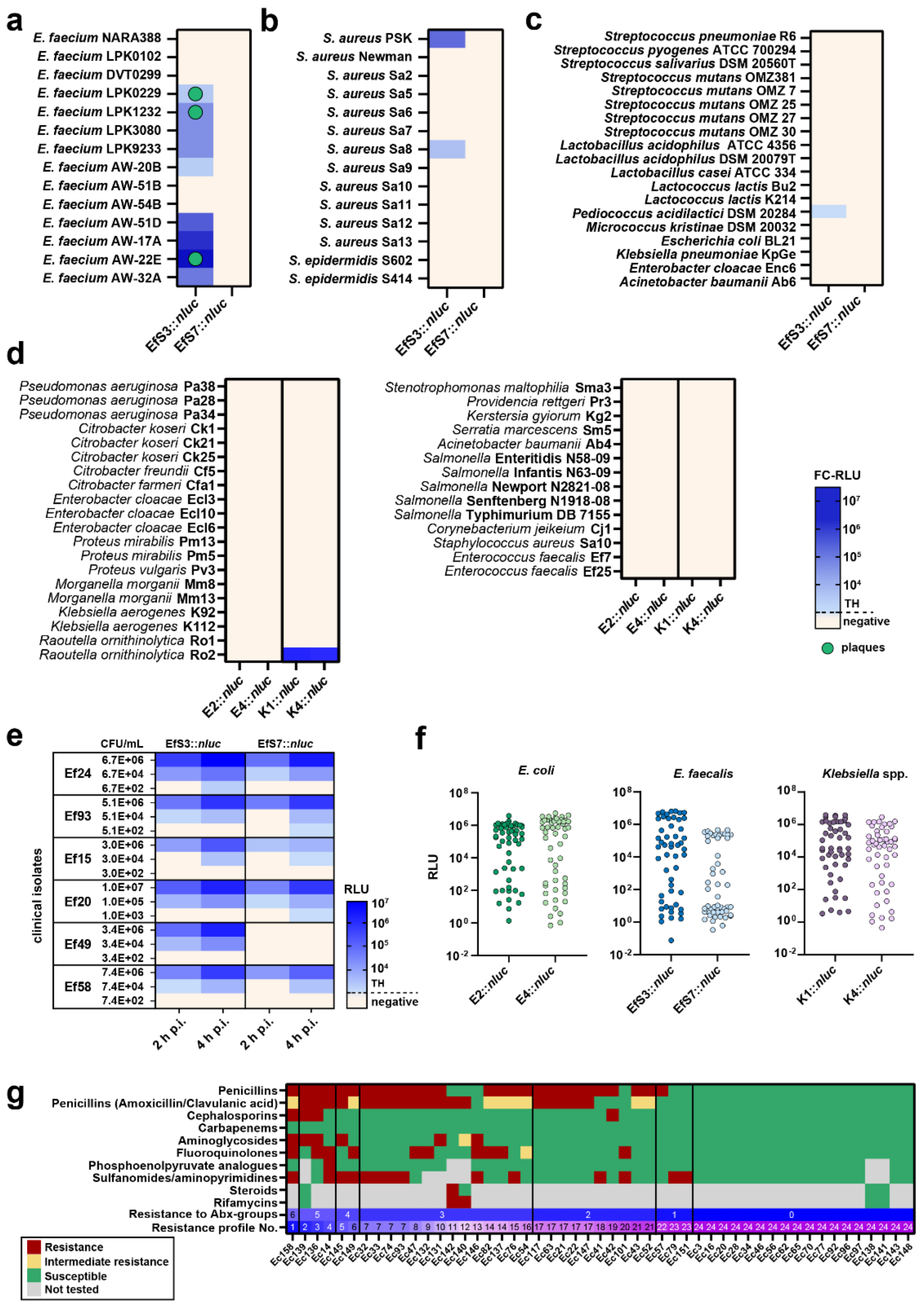

**Figure S3. Reporter phage specificity analysis, activity in blood, and clonal diversity within the Zurich Uropathogen Collection, related to Fig. 4.** To address phage specificity, the plaquing host ranges and bioluminescence detection ranges of phages EfS3*::nluc* and EfS7*::nluc* were determined on the indicated (**a**) *E. faecium* or (**b**) *S. aureus* strains and (**c**) on a set of additional Gram-positive and Gram-negative bacteria. (**d**) A similar specificity analysis was performed with reporter phages E2*::nluc*, E4*::nluc*, K1*::nluc*, and K4*::nluc*. The fold-change RLU (FC-RLU) is shown in shades of blue, plaque formation is indicated with a green dot. Thresholds (TH) are 50 in panel (a), (b) and (c) and 100 in panel (d). Values are mean from biological triplicates. (**e**) Heat map showing reporter phage-induced luminescence in human blood spiked with 6 selected strains of *E. faecalis*. After 1/10 dilution of the spiked blood with buffered media and a 1h enrichment, reactions were mixed with 2x10$^{7}$ PFU/mL *Enterococcus* reporter phages and RLU was determined at 2 h and 4 h p.i., TH is 10$^{3}$ RLU (EfS3) and 2.5x10$^{2}$ (EfS7) (**f-g**) Analysis of the clonal diversity within the strain panel selected for host-range analysis. (**f**) To demonstrate strain diversity with respect to phage-sensitivity, the luminescence values (FC-RLU) from each reporter phage – host pair were plotted to visualize differences in signal production. Values are mean from biological triplicates. (**g**) To demonstrate strain diversity with respect to antibiotic sensitivity, all *E. coli* strains from the host range analysis were clustered according to their antibiotic resistance profiles. From 51 tested strains, 24 distinct patterns were identified, suggesting that E. *coli* clonal diversity is high.

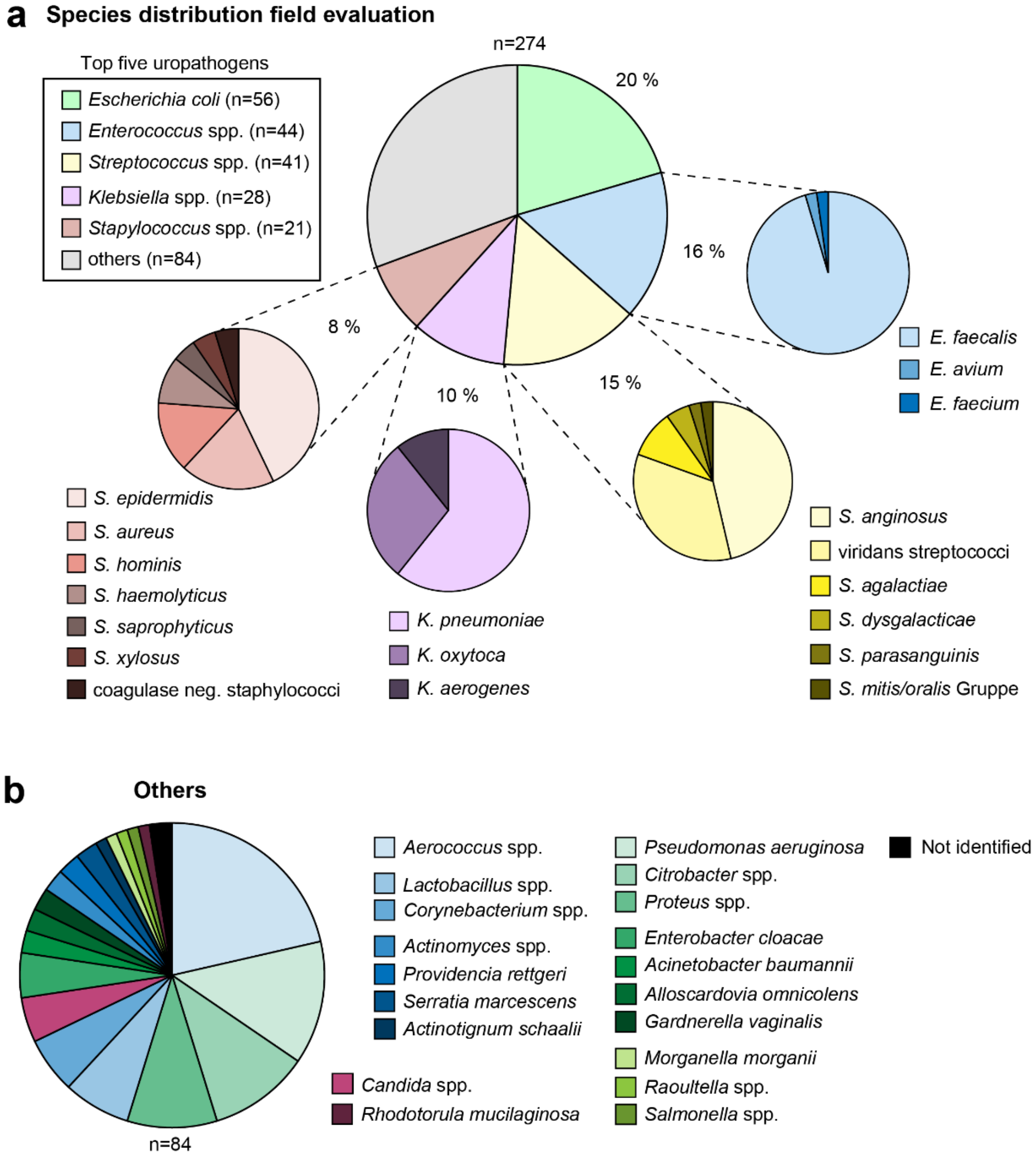


**Figure S4. Species distribution analysis of strains from the field evaluation, related to Fig. 5.** (**a**) A total of 274 bacterial stains identified during the field evaluation were analyzed with respect to their genus- and species distribution. Isolates that do not belong to the top five most prevalent uropathogens are shown in (**b**).

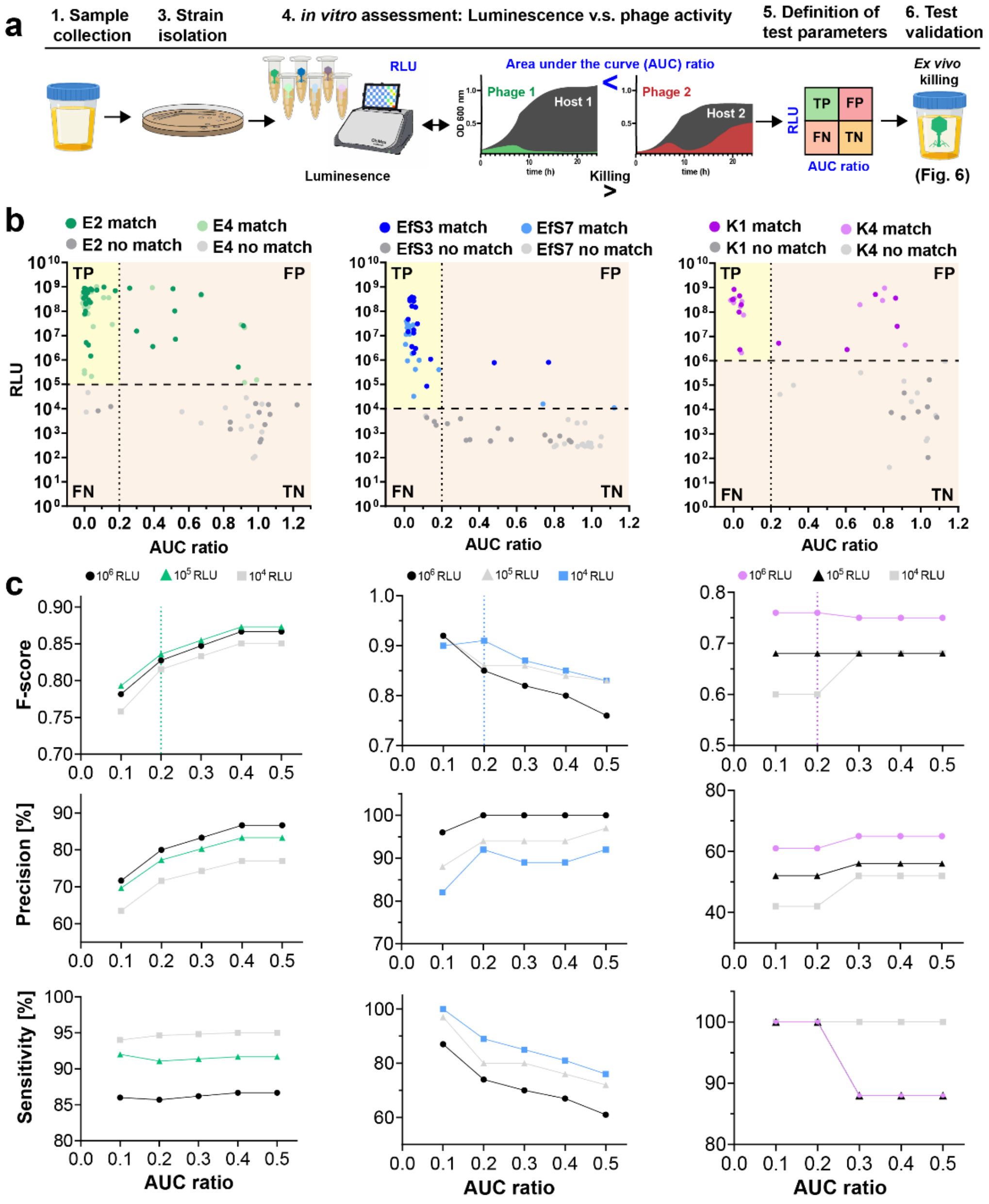


**Figure S5. Definition of test parameters for a reporter phage-based companion diagnostic, related to Fig. 6.** To develop a companion diagnostic assay that predicts phage antimicrobial activity, we correlated reporter phage-induced bioluminescence (RLU) with killing activity as determined by turbidity reduction assays (AUC ratio). Workflow shown in (**a**). The optimal RLU- and AUC ratio cutoffs were determined using defined experimental conditions. To this end, bacterial strains from the field evaluation were isolated and their luminescence response quantified using standardized infection conditions (starting phage titer = $10^7$ PFU / ml; starting bacterial concentration = OD0.01). Activity was quantified as the ratio of the area under the curve (AUC ratio: $AUC_{HOST+PHAGE}$ / $AUC_{HOST}$) where a value of one corresponds to no activity and a value of zero corresponds to maximal activity. (**b**) For each phage-host pair, the AUC ratio was plotted against the raw luminescence value. (**c**) Test parameters that produce the most reliable prediction of success while maintaining a stringent activity cut-off were identified by comparison of F1-score, precision, and sensitivity (see methods for more details).

### Supplementary Tables

**Table S1. The Zurich Uropathogen Collection.**

**Table S2. Strains used in this study.**

**Table S3. Bacteria isolated during the field evaluation from patient urine.**

**Table S4: Plasmids used and constructed during this study.**

**Table S5: Primer and synthetic DNA string sequences used in this study.**

**Table S1a.** ***E. coli*** **isolates of the Zurich Uropathogen Collection.**

| Species | Isolation month | UTI/ASB diagnosis | Isolate designation |
|---|---|---|---|
| *Escherichia coli* | January | ASB | Ec 1 |
| *Escherichia coli* | January | UTI | Ec 3 |
| *Escherichia coli* | January | ASB | Ec 4 |
| *Escherichia coli* | January | ASB | Ec 6 |
| *Escherichia coli* | January | ASB | Ec 10 |
| *Escherichia coli* | January | ASB | Ec 13 |
| *Escherichia coli* | January | ASB | Ec 12 |
| *Escherichia coli* | January | ASB | Ec 9 |
| *Escherichia coli* | January | UTI | Ec 11 |
| *Escherichia coli* | January | ASB | Ec 5 |
| *Escherichia coli* | January | ASB | Ec 2 |
| *Escherichia coli* | January | ASB | Ec 18 |
| *Escherichia coli* | January | ASB | Ec 17 |
| *Escherichia coli* | January | UTI | Ec 16 |
| *Escherichia coli* | January | ASB | Ec 15 |
| *Escherichia coli* | January | UTI | Ec 14 |
| *Escherichia coli* | January | ASB | Ec 8 |
| *Escherichia coli* | January | ASB | Ec 7 |
| *Escherichia coli* | January | ASB | Ec 19 |
| *Escherichia coli* | January | ASB | Ec 27 |
| *Escherichia coli* | January | ASB | Ec 26 |
| *Escherichia coli* | January | ASB | Ec 24 |
| *Escherichia coli* | January | ASB | Ec 23 |
| *Escherichia coli* | January | UTI | Ec 21 |
| *Escherichia coli* | January | ASB | Ec 168 |
| *Escherichia coli* | January | ASB | Ec 169 |
| *Escherichia coli* | January | UTI | Ec20 |
| *Escherichia coli* | January | UTI | Ec 22 |
| *Escherichia coli* | February | ASB | Ec 29 |
| *Escherichia coli* | February | ASB | Ec 30 |
| *Escherichia coli* | February | ASB | Ec 37 |
| *Escherichia coli* | February | ASB | Ec 39 |
| *Escherichia coli* | February | UTI | Ec 28 |
| *Escherichia coli* | February | ASB | Ec 38 |
| *Escherichia coli* | February | ASB | Ec 36 |
| *Escherichia coli* | February | UTI | Ec 32 |
| *Escherichia coli* | February | ASB | Ec 31 |
| *Escherichia coli* | February | UTI | Ec 33 |
| *Escherichia coli* | February | UTI | Ec 34 |
| *Escherichia coli* | February | ASB | Ec 35 |
| *Escherichia coli* | February | UTI | Ec 42 |
| *Escherichia coli* | February | UTI | Ec 46 |
| *Escherichia coli* | February | ASB | Ec 48 |
| *Escherichia coli* | February | UTI | Ec 47 |
| *Escherichia coli* | February | ASB | Ec 51 |
| *Escherichia coli* | February | ASB | Ec 40 |
| *Escherichia coli* | February | ASB | Ec 50 |
| *Escherichia coli* | February | UTI | Ec 41 |
| *Escherichia coli* | February | ASB | Ec 44 |
| *Escherichia coli* | February | UTI | Ec 52 |
| *Escherichia coli* | February | ASB | Ec 53 |
| *Escherichia coli* | February | ASB | Ec 45 |
| *Escherichia coli* | February | UTI | Ec 43 |
| *Escherichia coli* | February | ASB | Ec 61 |
| *Escherichia coli* | February | UTI | Ec 57 |
| *Escherichia coli* | February | UTI | Ec 54 |
| *Escherichia coli* | February | UTI | Ec 62 |
| *Escherichia coli* | February | UTI | Ec 63 |
| *Escherichia coli* | February | UTI | Ec 65 |
| *Escherichia coli* | February | ASB | Ec 55 |
| *Escherichia coli* | February | UTI | Ec 56 |
| *Escherichia coli* | February | ASB | Ec 58 |
| *Escherichia coli* | February | ASB | Ec 60 |
| *Escherichia coli* | February | ASB | Ec 64 |
| *Escherichia coli* | February | UTI | Ec 79 |
| *Escherichia coli* | February | ASB | n/a |
| *Escherichia coli* | February | UTI | Ec 77 |
| *Escherichia coli* | February | ASB | Ec 73 |
| *Escherichia coli* | February | UTI | Ec 76 |
| *Escherichia coli* | February | UTI | Ec 74 |
| *Escherichia coli* | February | UTI | Ec 80 |
| *Escherichia coli* | February | ASB | Ec 71 |
| *Escherichia coli* | February | ASB | Ec 78 |
| *Escherichia coli* | February | UTI | Ec 70 |
| *Escherichia coli* | February | ASB | Ec 72 |
| *Escherichia coli* | February | ASB | Ec 67 |
| *Escherichia coli* | February | ASB | Ec 66 |
| *Escherichia coli* | February | ASB | Ec 68 |
| *Escherichia coli* | February | ASB | Ec 69 |
| *Escherichia coli* | March | ASB | Ec 90 |
| *Escherichia coli* | March | ASB | Ec 94 |
| *Escherichia coli* | March | UTI | Ec 92 |
| *Escherichia coli* | March | UTI | Ec 93 |
| *Escherichia coli* | March | ASB | Ec 87 |
| *Escherichia coli* | March | ASB | Ec 88 |
| *Escherichia coli* | March | ASB | Ec 89 |
| *Escherichia coli* | March | UTI | Ec 96 |
| *Escherichia coli* | March | ASB | Ec 95 |
| *Escherichia coli* | March | UTI | Ec 97 |
| *Escherichia coli* | March | UTI | Ec 82 |
| *Escherichia coli* | March | ASB | Ec 91 |
| *Escherichia coli* | March | ASB | Ec222 |
| *Escherichia coli* | March | ASB | Ec 84 |
| *Escherichia coli* | March | UTI | Ec 83 |
| *Escherichia coli* | March | ASB | Ec 85 |
| *Escherichia coli* | March | UTI | Ec 86 |
| *Escherichia coli* | March | ASB | Ec 107 |
| *Escherichia coli* | March | ASB | Ec 103 |
| *Escherichia coli* | March | ASB | Ec 102 |
| *Escherichia coli* | March | UTI | Ec 104 |
| *Escherichia coli* | March | ASB | Ec 105 |
| *Escherichia coli* | March | ASB | Ec 100 |
| *Escherichia coli* | March | ASB | Ec 99 |
| *Escherichia coli* | March | ASB | Ec 98 |
| *Escherichia coli* | March | ASB | Ec 108 |
| *Escherichia coli* | March | ASB | Ec 106 |
| *Escherichia coli* | March | ASB | Ec 109 |
| *Escherichia coli* | March | ASB | Ec 110 |
| *Escherichia coli* | March | UTI | Ec 101 |
| *Escherichia coli* | March | ASB | Ec 135 |

**Table S1b.** ***Klebsiella*** **spp. isolates of the Zurich Uropathogen Collection.**

| Species | Isolation month | UTI/ASB diagnosis | Isolate designation |
|---|---|---|---|
| *Klebsiella oxytoca* | January | ASB | Ko 2 |
| *Klebsiella pneumoniae* | January | UTI | Kp 3 |
| *Klebsiella pneumoniae* | January | ASB | Kp 8 |
| *Klebsiella oxytoca* | January | ASB | Ko 7 |
| *Klebsiella pneumoniae* | January | ASB | Kp 6 |
| *Klebsiella pneumoniae* | January | ASB | Kp14 |
| *Klebsiella pneumoniae* | January | ASB | Kp13 |
| *Klebsiella pneumoniae* | January | ASB | Kp 11 |
| *Klebsiella pneumoniae* | January | UTI | Kp 106 |
| *Klebsiella pneumoniae* | January | UTI | Kp 5 |
| *Klebsiella pneumoniae* | January | ASB | Kp 15 |
| *Klebsiella pneumoniae* | January | ASB | Kp 9 |
| *Klebsiella pneumoniae* | January | ASB | Kp 17 |
| *Klebsiella pneumoniae* | January | ASB | Kp 18 |
| *Klebsiella pneumoniae* | January | ASB | Kp 20 |
| *Klebsiella pneumoniae* | January | UTI | Kp 16 |
| *Klebsiella pneumoniae* | January | ASB | Kp 19 |
| *Klebsiella pneumoniae* | February | ASB | Kp 25 |
| *Klebsiella pneumoniae* | February | ASB | Kp 24 |
| *Klebsiella pneumoniae* | February | ASB | Kp 23 |
| *Klebsiella pneumoniae* | February | ASB | Kp 31 |
| *Klebsiella pneumoniae* | February | ASB | Kp 30 |
| *Klebsiella pneumoniae* | February | ASB | Kp 28 |
| *Klebsiella pneumoniae* | February | UTI | Kp 26 |
| *Klebsiella pneumoniae* | February | ASB | Kp 27 |
| *Klebsiella oxytoca* | February | ASB | Ko 29 |
| *Klebsiella pneumoniae* | February | ASB | Kp 22 |
| *Klebsiella pneumoniae* | February | ASB | Kp 21 |
| *Klebsiella pneumoniae* | February | UTI | Kp 36 |
| *Klebsiella pneumoniae* | February | UTI | Kp 37 |
| *Klebsiella oxytoca* | February | UTI | Ko 38 |
| *Klebsiella pneumoniae* | February | UTI | Kp 34 |
| *Klebsiella pneumoniae* | February | ASB | Kp 32 |
| *Klebsiella pneumoniae* | February | ASB | Kp 41 |
| *Klebsiella pneumoniae* | February | ASB | Kp 35 |
| *Klebsiella pneumoniae* | February | ASB | Kp 40 |
| *Klebsiella pneumoniae* | February | UTI | Kp 39 |
| *Klebsiella pneumoniae* | February | UTI | Kp 43 |
| *Klebsiella pneumoniae* | February | UTI | Kp 48 |
| *Klebsiella oxytoca* | February | UTI | n/a |
| *Klebsiella pneumoniae* | February | UTI | Kp 45 |
| *Klebsiella pneumoniae* | February | ASB | Kp 50 |
| *Klebsiella pneumoniae* | February | ASB | Kp 52 |
| *Klebsiella oxytoca* | February | ASB | Ko 49 |
| *Klebsiella (Enterobacter) aerogenes* | February | ASB | Ka 44 |
| *Klebsiella pneumoniae* | February | UTI | Kp 51 |
| *Klebsiella pneumoniae* | February | ASB | Kp 46 |
| *Klebsiella pneumoniae* | February | ASB | Kp 47 |
| *Klebsiella pneumoniae* | March | ASB | Kp 61 |
| *Klebsiella pneumoniae* | March | UTI | Kp 66 |
| *Klebsiella pneumoniae* | March | ASB | Kp 60 |
| *Klebsiella oxytoca* | March | ASB | Ko 63 |
| *Klebsiella pneumoniae* | March | ASB | Kp 62 |
| *Klebsiella pneumoniae* | March | ASB | Kp 55 |
| *Klebsiella pneumoniae* | March | ASB | Kp 54 |
| *Klebsiella pneumoniae* | March | ASB | Kp 59 |
| *Klebsiella pneumoniae* | March | ASB | Kp 57 |
| *Klebsiella pneumoniae* | March | ASB | Kp 56 |
| *Klebsiella pneumoniae* | March | ASB | Kp 58 |
| *Klebsiella pneumoniae* | March | ASB | Kp 64 |
| *Klebsiella (Enterobacter) aerogenes* | March | ASB | Ka 65 |
| *Klebsiella pneumoniae* | March | ASB | Kp 72 |
| *Klebsiella pneumoniae* | March | ASB | Kp 70 |
| *Klebsiella pneumoniae* | March | ASB | Kp 71 |
| *Klebsiella pneumoniae* | March | ASB | Kp 74 |
| *Klebsiella pneumoniae* | March | ASB | Kp 73 |
| *Klebsiella oxytoca* | March | ASB | Ko 75 |
| *Klebsiella oxytoca* | March | UTI | Ko 68 |
| *Klebsiella pneumoniae* | March | UTI | Kp 69 |
| *Klebsiella pneumoniae* | March | ASB | Kp 67 |
| *Klebsiella pneumoniae* | March | ASB | Kp 82 |
| *Klebsiella pneumoniae* | March | ASB | Kp 83 |
| *Klebsiella oxytoca* | March | ASB | Ko 84 |
| *Klebsiella pneumoniae* | March | ASB | Kp 85 |
| *Klebsiella pneumoniae* | March | ASB | Kp 80 |
| *Klebsiella pneumoniae* | March | ASB | Kp 81 |
| *Klebsiella pneumoniae* | April | ASB | Kp 76 |
| *Klebsiella oxytoca* | April | ASB | Ko 77 |
| *Klebsiella pneumoniae* | April | ASB | Kp 78 |
| *Klebsiella pneumoniae* | April | ASB | Kp 79 |
| *Klebsiella oxytoca* | April | UTI | Ko 87 |
| *Klebsiella pneumoniae* | April | UTI | Kp 93 |
| *Klebsiella pneumoniae* | April | UTI | Kp 90 |
| *Klebsiella pneumoniae* | April | UTI | Kp 89 |
| *Klebsiella pneumoniae* | April | UTI | Kp 88 |
| *Klebsiella oxytoca* | April | UTI | Ko 91 |
| *Klebsiella (Enterobacter) aerogenes* | April | UTI | Ka 92 |
| *Klebsiella pneumoniae* | May | UTI | Kp 94 |
| *Klebsiella pneumoniae* | May | UTI | Kp 98 |
| *Klebsiella (Enterobacter) aerogenes* | May | UTI | Ka 99 |
| *klebsiella pneumoniae* | May | UTI | Kp 96 |
| *Klebsiella pneumoniae* | May | UTI | Kp 95 |
| *Klebsiella pneumoniae* | May | UTI | Kp 97 |
| *Klebsiella oxytoca* | May | UTI | Ko 101 |
| *Klebsiella pneumoniae* | May | UTI | Kp 100 |
| *Klebsiella pneumoniae* | June | UTI | Kp 102 |
| *Klebsiella pneumoniae* | June | UTI | Kp 103 |
| *Klebsiella pneumoniae* | July | UTI | Kp 104 |
| *Klebsiella pneumoniae* | July | UTI | Kp 105 |
| *Klebsiella pneumoniae* | August | UTI | Kp 109 |
| *Klebsiella pneumoniae* | August | UTI | Kp 108 |
| *Klebsiella pneumoniae* | August | UTI | Kp 107 |
| *Klebsiella pneumoniae* | August | UTI | Kp 110 |
| *Klebsiella (Enterobacter) aerogenes* | August | UTI | Ka 114 |
| *Klebsiella pneumoniae* | August | UTI | Kp 111 |
| *Klebsiella (Enterobacter) aerogenes* | August | UTI | Ka 112 |
| *Klebsiella pneumoniae* | September | UTI | Kp 113 |
| *Klebsiella (Enterobacter) aerogenes* | September | UTI | Ka 119 |
| *Klebsiella (Enterobacter) aerogenes* | September | UTI | Ka 115 |
| *Klebsiella pneumoniae* | September | UTI | Kp 118 |

| | | | |
|---|---|---|---|
| *Escherichia coli* | March | ASB | Ec 126 |
| *Escherichia coli* | March | ASB | Ec 128 |
| *Escherichia coli* | March | UTI | Ec 129 |
| *Escherichia coli* | March | ASB | Ec 130 |
| *Escherichia coli* | March | UTI | Ec 131 |
| *Escherichia coli* | April | UTI | Ec 132 |
| *Escherichia coli* | April | ASB | Ec 133 |
| *Escherichia coli* | April | UTI | Ec 134 |
| *Escherichia coli* | April | ASB | Ec 111 |
| *Escherichia coli* | April | ASB | Ec 112 |
| *Escherichia coli* | April | ASB | Ec 113 |
| *Escherichia coli* | April | ASB | Ec 114 |
| *Escherichia coli* | April | ASB | Ec 115 |
| *Escherichia coli* | April | ASB | Ec 116 |
| *Escherichia coli* | April | UTI | Ec 117 |
| *Escherichia coli* | April | ASB | Ec 118 |
| *Escherichia coli* | April | ASB | Ec 119 |
| *Escherichia coli* | April | ASB | Ec 120 |
| *Escherichia coli* | April | ASB | Ec 121 |
| *Escherichia coli* | April | ASB | Ec 122 |
| *Escherichia coli* | April | ASB | Ec 123 |
| *Escherichia coli* | April | ASB | Ec 124 |
| *Escherichia coli* | April | ASB | Ec 125 |
| *Escherichia coli* | April | ASB | n/a |
| *Escherichia coli* | April | UTI | Ec 136 |
| *Escherichia coli* | April | UTI | Ec 138 |
| *Escherichia coli* | April | UTI | Ec 142 |
| *Escherichia coli* | April | UTI | Ec 137 |
| *Escherichia coli* | April | UTI | Ec 139 |
| *Escherichia coli* | April | UTI | Ec 141 |
| *Escherichia coli* | May | UTI | Ec 140 |
| *Escherichia coli* | May | UTI | Ec 143 |
| *Escherichia coli* | May | UTI | Ec 145 |
| *Escherichia coli* | May | UTI | Ec 146 |
| *Escherichia coli* | May | UTI | Ec 147 |
| *Escherichia coli* | May | UTI | Ec 148 |
| *Escherichia coli* | May | UTI | Ec 149 |
| *Escherichia coli* | May | UTI | Ec 151 |
| *Escherichia coli* | May | UTI | Ec 150 |
| *Escherichia coli* | May | UTI | Ec 153 |
| *Escherichia coli* | May | UTI | Ec 152 |
| *Escherichia coli* | June | UTI | Ec 154 |
| *Escherichia coli* | June | UTI | Ec 155 |
| *Escherichia coli* | June | UTI | Ec 156 |
| *Escherichia coli* | June | UTI | Ec 158 |
| *Escherichia coli* | June | UTI | Ec 159 |
| *Escherichia coli* | June | UTI | Ec 157 |
| *Escherichia coli* | July | UTI | Ec 160 |
| *Escherichia coli* | July | UTI | Ec 161 |
| *Escherichia coli* | July | UTI | Ec 162 |
| *Escherichia coli* | July | UTI | Ec 167 |
| *Escherichia coli* | July | UTI | Ec 165 |
| *Escherichia coli* | July | UTI | Ec 166 |
| *Escherichia coli* | August | ASB | Ec 171 |
| *Escherichia coli* | August | UTI | Ec 170 |
| *Escherichia coli* | August | UTI | Ec 164 |
| *Escherichia coli* | August | UTI | Ec 163 |
| *Escherichia coli* | August | UTI | Ec 173 |
| *Escherichia coli* | August | UTI | Ec 174 |
| *Escherichia coli* | August | UTI | Ec 172 |
| *Escherichia coli* | August | UTI | Ec 178 |
| *Escherichia coli* | August | UTI | Ec 175 |
| *Escherichia coli* | September | UTI | Ec 176 |
| *Escherichia coli* | September | UTI | Ec 177 |
| *Escherichia coli* | September | UTI | Ec 179 |
| *Escherichia coli* | September | UTI | Ec 182 |
| *Escherichia coli* | October | UTI | Ec 180 |
| *Escherichia coli* | October | UTI | Ec 181 |
| *Escherichia coli* | October | UTI | Ec 184 |
| *Escherichia coli* | October | UTI | Ec 185 |
| *Escherichia coli* | October | UTI | Ec 186 |
| *Escherichia coli* | October | UTI | Ec 187 |
| *Escherichia coli* | October | UTI | Ec 188 |
| *Escherichia coli* | October | UTI | Ec 189 |
| *Escherichia coli* | October | UTI | Ec 190 |
| *Escherichia coli* | October | UTI | Ec 192 |
| *Escherichia coli* | October | UTI | Ec 193 |
| *Escherichia coli* | November | UTI | Ec 194 |
| *Escherichia coli* | November | UTI | Ec 195 |
| *Escherichia coli* | November | UTI | Ec 198 |
| *Escherichia coli* | November | UTI | Ec 200 |
| *Escherichia coli* | November | UTI | Ec 202 |
| *Escherichia coli* | November | UTI | Ec 204 |
| *Escherichia coli* | November | UTI | Ec 210 |
| *Escherichia coli* | November | UTI | Ec 208 |
| *Escherichia coli* | November | UTI | Ec 209 |
| *Escherichia coli* | November | UTI | Ec 211 |
| *Escherichia coli* | November | UTI | Ec 212 |
| *Escherichia coli* | November | UTI | Ec 221 |
| *Escherichia coli* | November | UTI | Ec 213 |
| *Escherichia coli* | December | UTI | Ec 214 |
| *Escherichia coli* | December | UTI | Ec 215 |
| *Escherichia coli* | December | UTI | Ec 216 |
| *Escherichia coli* | December | UTI | Ec 217 |
| *Escherichia coli* | December | UTI | Ec 218 |
| *Escherichia coli* | December | UTI | Ec 219 |
| *Escherichia coli* | December | UTI | Ec 220 |

| | | | |
|---|---|---|---|
| *Klebsiella pneumoniae* | September | UTI | Kp 120 |
| *Klebsiella pneumoniae* | October | UTI | Kp 116 |
| *Klebsiella pneumoniae* | October | UTI | Kp 117 |
| *Klebsiella pneumoniae* | October | UTI | Kp 121 |
| *Klebsiella pneumoniae* | October | UTI | Kp 112 |
| *Klebsiella pneumoniae* | October | UTI | Kp 123 |
| *Klebsiella pneumoniae* | October | UTI | Kp 124 |
| *Klebsiella pneumoniae* | October | UTI | Kp 125 |
| *Klebsiella pneumoniae* | November | UTI | Kp 126 |
| *Klebsiella pneumoniae* | November | UTI | Kp 127 |

**Table S1c. *Enterococcus* spp. isolates of the Zurich Uropathogen Collection.**

| Species | Isolation month | UTI/ASB diagnosis | Isolate designation |
|---|---|---|---|
| *Enterococcus faecalis* | January | ASB | Ef 8 |
| *Enterococcus faecalis* | January | UTI | Ef 4 |
| *Enterococcus faecalis* | January | ASB | Ef 1 |
| *Enterococcus faecalis* | January | UTI | Ef 2 |
| *Enterococcus faecalis* | January | ASB | Ef 3 |
| *Enterococcus faecalis* | January | ASB | Ef 6 |
| *Enterococcus faecalis* | January | ASB | Ef 5 |
| *Enterococcus faecalis* | January | UTI | Ef 7 |
| *Enterococcus faecalis* | February | ASB | Ef 15 |
| *Enterococcus faecalis* | February | ASB | Ef 16 |
| *Enterococcus faecalis* | February | ASB | Ef 18 |
| *Enterococcus faecalis* | February | UTI | Ef 9 |
| *Enterococcus faecalis* | February | ASB | Ef 10 |
| *Enterococcus faecalis* | February | ASB | Ef 17 |
| *Enterococcus faecalis* | February | UTI | Ec 12 |
| *Enterococcus faecalis* | February | ASB | Ef 21 |
| *Enterococcus avium* | February | UTI | Ea 1 |
| *Enterococcus faecalis* | February | UTI | Ef 20 |
| *Enterococcus faecalis* | February | ASB | Ef 11 |
| *Enterococcus faecalis* | February | ASB | Ef 13 |
| *Enterococcus faecalis* | February | ASB | Ef 14 |
| *Enterococcus faecalis* | February | ASB | Ef 19 |
| *Enterococcus faecalis* | February | ASB | Ef 24 |
| *Enterococcus faecalis* | February | UTI | Ef 25 |
| *Enterococcus faecalis* | February | ASB | Ef 28 |
| *Enterococcus faecalis* | February | UTI | Ef 27 |
| *Enterococcus faecalis* | February | ASB | Ef 22 |
| *Enterococcus faecalis* | February | UTI | Ef 26 |
| *Enterococcus faecalis* | February | ASB | Ef 23 |
| *Enterococcus faecalis* | February | ASB | Ef 32 |
| *Enterococcus faecalis* | February | UTI | Ef 34 |
| *Enterococcus faecalis* | February | ASB | Ef 31 |
| *Enterococcus faecalis* | February | UTI | n/a |
| *Enterococcus faecalis* | February | UTI | Ef 30 |
| *Enterococcus faecalis* | February | ASB | Ef 29 |
| *Enterococcus faecalis* | February | ASB | Ef 33 |
| *Enterococcus faecalis* | February | ASB | Ef 37 |
| *Enterococcus faecalis* | February | UTI | Ef 36 |
| *Enterococcus faecalis* | February | ASB | Ef 42 |
| *Enterococcus faecalis* | February | UTI | Ef 40 |
| *Enterococcus faecalis* | February | ASB | Ef 43 |
| *Enterococcus faecalis* | February | UTI | Ef 48 |
| *Enterococcus faecalis* | February | UTI | Ef 47 |
| *Enterococcus faecalis* | February | ASB | Ef 45 |
| *Enterococcus faecalis* | February | ASB | Ef 44 |
| *Enterococcus faecalis* | February | ASB | Ef 39 |
| *Enterococcus faecalis* | February | ASB | Ef 46 |
| *Enterococcus faecalis* | February | ASB | Ef 38 |
| *Enterococcus faecalis* | March | ASB | Ef 61 |
| *Enterococcus faecalis* | March | UTI | Ef 59 |
| *Enterococcus faecalis* | March | UTI | Ef 57 |
| *Enterococcus faecalis* | March | ASB | Ef 53 |
| *Enterococcus faecalis* | March | ASB | Ef 60 |
| *Enterococcus faecalis* | March | ASB | Ef 52 |
| *Enterococcus faecalis* | March | UTI | Ef 58 |
| *Enterococcus faecalis* | March | ASB | Ef 50 |
| *Enterococcus faecalis* | March | ASB | Ef 49 |
| *Enterococcus faecalis* | March | ASB | Ef 51 |
| *Enterococcus faecalis* | March | ASB | Ef 62 |
| *Enterococcus faecalis* | March | ASB | Ef 54 |
| *Enterococcus faecalis* | March | ASB | Ef 55 |
| *Enterococcus faecalis* | March | ASB | n/a |
| *Enterococcus faecalis* | March | ASB | n/a |
| *Enterococcus faecalis* | March | ASB | n/a |
| *Enterococcus faecalis* | March | ASB | Ef 64 |
| *Enterococcus faecalis* | March | ASB | n/a |
| *Enterococcus faecalis* | March | ASB | Ef 63 |
| *Enterococcus faecalis* | March | ASB | Ef 66 |
| *Enterococcus faecalis* | March | UTI | Ef 65 |
| *Enterococcus faecalis* | March | UTI | Ef 81 |
| *Enterococcus faecalis* | March | ASB | Ef 82 |
| *Enterococcus faecalis* | March | ASB | Ef 83 |
| *Enterococcus faecalis* | March | ASB | Ef 84 |
| *Enterococcus faecalis* | March | ASB | Ef85 |
| *Enterococcus faecalis* | March | UTI | Ef 74 |
| *Enterococcus faecalis* | March | ASB | Ef 76 |
| *Enterococcus faecalis* | March | ASB | Ef 77 |
| *Enterococcus faecium* | March | ASB | Efm 1 |
| *Enterococcus faecalis* | March | ASB | Ef 78 |
| *Enterococcus faecalis* | April | ASB | Ef 79 |
| *Enterococcus faecalis* | April | ASB | Ef 80 |
| *Enterococcus faecalis* | April | ASB | Ef67 |
| *Enterococcus faecalis* | April | ASB | Ef 68 |
| *Enterococcus faecalis* | April | ASB | Ef 69 |
| *Enterococcus faecalis* | April | ASB | Ef 70 |
| *Enterococcus faecalis* | April | ASB | Ef 71 |
| *Enterococcus faecalis* | April | UTI | Ef 72 |
| *Enterococcus faecalis* | April | ASB | Ef 73 |
| *Enterococcus faecalis* | April | ASB | Ef 75 |
| *Enterococcus faecalis* | April | UTI | Ef 90 |
| *Enterococcus faecalis* | April | UTI | Ef 89 |
| *Enterococcus faecalis* | April | UTI | Ef 91 |
| *Enterococcus faecalis* | April | UTI | Ef 88 |
| *Enterococcus faecalis* | May | UTI | Ef 94 |
| *Enterococcus faecalis* | May | UTI | Ef 92 |
| *Enterococcus faecalis* | May | UTI | Ef 93 |
| *Enterococcus faecalis* | May | UTI | Ef 96 |
| *Enterococcus faecalis* | June | UTI | Ef 95 |
| *Enterococcus faecalis* | July | UTI | Ef 97 |
| *Enterococcus faecalis* | July | UTI | Ef 98 |
| *Enterococcus faecalis* | July | UTI | Ef 99 |
| *Enterococcus faecalis* | July | UTI | Ef 100 |
| *Enterococcus faecalis* | July | UTI | Ef 103 |
| *Enterococcus faecalis* | August | UTI | Ef 109 |
| *Enterococcus faecalis* | August | UTI | Ef 102 |
| *Enterococcus faecalis* | August | UTI | Ef 101 |
| *Enterococcus faecalis* | August | UTI | Ef 104 |
| *Enterococcus faecalis* | August | UTI | Ef 105 |
| *Enterococcus faecalis* | August | UTI | Ef 106 |
| *Enterococcus faecalis* | August | UTI | Ef 107 |
| *Enterococcus faecalis* | September | UTI | Ef 108 |
| *Enterococcus faecalis* | September | UTI | Ef 110 |
| *Enterococcus faecalis* | September | UTI | Ef 112 |
| *Enterococcus faecalis* | October | UTI | Ef 111 |
| *Enterococcus avium* | October | UTI | Ea 2 |
| *Enterococcus faecalis* | October | UTI | Ef 113 |
| *Enterococcus faecalis* | October | UTI | Ef 114 |
| *Enterococcus faecalis* | October | UTI | Ef 115 |
| *Enterococcus faecalis* | October | UTI | Ef 116 |

**Table S1d. Other species isolated as part of the Zurich Uropathogen Collection.**

| Species | Isolation month | UTI/ASB diagnosis | Isolate designation |
|---|---|---|---|
| *Citrobacter koseri (diversus)* | January | ASB | Ck 1 |
| *Pseudomonas aeruginosa* | January | ASB | Pa 1 |
| *Actinotignum (Actinobaculum) schaalii* | January | ASB | n/a |
| *Acinetobacter baumannii* | January | UTI | Ab 4 |
| *Enterobacter cloacae* | January | UTI | Enc 10 |
| *Proteus vulgaris/penneri* | January | ASB | Pvp 1 |
| *Pseudomonas aeruginosa* | January | ASB | Pa 3 |
| *Pseudomonas aeruginosa* | January | ASB | Pa 2 |
| *Pseudomonas aeruginosa* | January | ASB | Pa 38 |
| *Stenotrophomonas maltophilia* | February | ASB | Sm 1 |
| *Staphylococcus aureus* | February | ASB | Sa 1 |
| *Proteus mirabilis* | February | UTI | Pm 1 |
| *Pseudomonas aeruginosa* | February | ASB | Pa 4 |
| *Proteus mirabilis* | February | ASB | Pm 3 |
| *Proteus vulgaris* | February | ASB | Pv 1 |
| *Proteus mirabilis* | February | UTI | Pm 2 |
| *Acinetobacter baumannii Komplex* | February | ASB | Ab 1 |
| *Citrobacter koseri (diversus)* | February | ASB | Ck 2 |
| *Pseudomonas aeruginosa* | February | ASB | Pa 5 |
| *Pseudomonas aeruginosa* | February | ASB | Pa 7 |
| *Morganella morganii* | February | ASB | Mm 2 |
| *Proteus mirabilis* | February | UTI | Pm 4 |
| *Morganella morganii* | February | ASB | Mm 1 |
| *Pseudomonas aeruginosa* | February | UTI | Pa 8 |
| *Citrobacter koseri (diversus)* | February | ASB | Ck 4 |
| *Proteus mirabilis* | February | UTI | Pm 6 |
| *Pseudomonas aeruginosa* | February | ASB | Pa 6 |
| *Citrobacter freundii* | February | ASB | Cf 1 |
| *Pseudomonas aeruginosa* | February | ASB | Pa 9 |
| *Citrobacter freundii* | February | ASB | Cf 2 |
| *Proteus mirabilis* | February | UTI | Pm 5 |
| *Citrobacter koseri (diversus)* | February | ASB | Ck 3 |
| *Serratia marcescens* | February | ASB | Sms 1 |
| *Pseudomonas aeruginosa* | February | ASB | Pa 10 |
| *Citrobacter koseri (diversus)* | February | ASB | Ck 6 |
| *Citrobacter koseri (diversus)* | February | ASB | Ck 5 |
| *Serratia marcescens* | February | ASB | Sms 2 |
| *Proteus mirabilis* | February | ASB | Pm 7 |
| *Proteus vulgaris/penneri* | February | ASB | Pvp 2 |
| *Enterobacter cloacae* | February | UTI | Enc 1.1 |
| *Enterobacter cloacae* | February | ASB | Enc 2.1 |
| *Staphylococcus aureus* | February | ASB | Sms 3 |
| *Serratia marcescens* | February | ASB | Sa 3 |
| *Pseudomonas aeruginosa* | February | UTI | Pa 13 |
| *Proteus vulgaris/penneri* | February | UTI | Pvp 3 |
| *Staphylococcus aureus* | February | ASB | Sa 4 |
| *Citrobacter koseri (diversus)* | February | ASB | Ck 9 |
| *Proteus mirabilis* | February | ASB | Pm 8 |
| *Proteus mirabilis* | February | UTI | Pm 9 |
| *Pseudomonas aeruginosa* | February | UTI | Pa 12 |
| *Acinetobacter baumannii* | February | UTI | Ac 2 |
| *Staphylococcus aureus* | February | ASB | Sa 2 |
| *Enterobacter cloacae* | February | UTI | Enc 3.1 |
| *Morganella morganii* | February | UTI | Mm 4 |
| *Morganella morganii* | February | ASB | Mm 3 |
| *Citrobacter koseri (diversus)* | February | ASB | Ck 10 |
| *Pseudomonas aeruginosa* | February | UTI | Pa 11 |
| *Citrobacter koseri (diversus)* | February | ASB | Ck 7 |
| *Citrobacter koseri (diversus)* | February | ASB | Ck 8 |
| *Proteus mirabilis* | March | ASB | Pm 11 |
| *Staphylococcus aureus* | March | ASB | Sa 5 |
| *Actinotignum (Actinobaculum) schaalii* | March | UTI | As 1 |
| *Pseudomonas aeruginosa* | March | ASB | Pa 16 |
| *Providencia rettgeri* | March | ASB | Pr 1 |
| *Proteus vulgaris/penneri* | March | ASB | Pvp 6 |
| *Citrobacter koseri (diversus)* | March | ASB | Ck 11 |
| *Pseudomonas aeruginosa* | March | ASB | Pa 18 |
| *Staphylococcus aureus* | March | ASB | Sa 6 |
| *Morganella morganii* | March | ASB | Mm 6 |
| *Raoutella (Klebsiella) ornithinolytica* | March | ASB | Ro 1 |
| *Pseudomonas aeruginosa* | March | ASB | Pa 17 |
| *Citrobacter koseri (diversus)* | March | ASB | Ck 13 |
| *Citrobacter freundii* | March | ASB | Cf 3 |
| *Staphylococcus aureus* | March | ASB | Sa 7 |
| *Proteus vulgaris/penneri* | March | ASB | Pvp 5 |
| *Pseudomonas aeruginosa* | March | ASB | Pa 15 |
| *Pseudomonas aeruginosa* | March | ASB | Pa 19 |
| *Morganella morganii* | March | ASB | Mm 5 |
| *Citrobacter koseri (diversus)* | March | ASB | Ck 14 |
| *Pseudomonas aeruginosa* | March | ASB | Pa 14 |
| *Citrobacter koseri (diversus)* | March | ASB | Ck 12 |
| *Actinotignum (Actinobaculum) schaalii* | March | ASB | As 2 |
| *Proteus mirabilis* | March | ASB | Pm 10 |
| *Morganella morganii* | March | ASB | Mm 7 |
| *Morganella morganii* | March | ASB | Mm 8 |
| *Citrobacter koseri (diversus)* | March | ASB | Ck 15 |
| *Morganella morganii* | March | ASB | Mm 10 |
| *Pseudomonas aeruginosa* | March | ASB | Pa 23 |
| *Citrobacter koseri (diversus)* | March | ASB | Ck 17 |
| *Staphylococcus aureus* | March | ASB | Sa 8 |
| *Citrobacter koseri (diversus)* | March | ASB | Ck 16 |
| *Pseudomonas aeruginosa* | March | ASB | Pa 24 |
| *Raoutella (Klebsiella) ornithinolytica* | March | ASB | Ro 2 |
| *Serratia marcescens* | March | UTI | Sms 5 |
| *Morganella morganii* | March | UTI | Mm 9 |
| *Pseudomonas aeruginosa* | March | ASB | Pa 21 |
| *Pseudomonas aeruginosa* | March | ASB | Pa 22 |
| *Pseudomonas aeruginosa* | March | ASB | Pa 20 |
| *Serratia marcescens* | March | ASB | Sms 4 |
| *Serratia marcescens* | March | ASB | Sms 7 |
| *Pseudomonas aeruginosa* | March | ASB | Pa 31 |
| *Proteus vulgaris/penneri* | March | ASB | Pvp 9 |
| *Staphylococcus aureus* | March | ASB | Sa 10 |
| *Proteus mirabilis* | March | ASB | Pm 13 |
| *Actinotignum (Actinobaculum) schaalii* | March | ASB | As 3 |
| *Citrobacter koseri (diversus)* | March | ASB | Ck 21 |
| *Pseudomonas aeruginosa* | March | ASB | Pa 28 |
| *Pseudomonas aeruginosa* | March | ASB | Pa 29 |
| *Pseudomonas aeruginosa* | March | UTI | Pa 30 |
| *Acinetobacter baumannii Komplex* | April | ASB | Ab 3 |
| *Stenotrophomonas maltophilia* | April | ASB | Sm 2 |
| *Citrobacter koseri (diversus)* | April | UTI | Ck 18 |
| *Pseudomonas aeruginosa* | April | ASB | Pa 25 |
| *Serratia marcescens* | April | ASB | Sms 6 |
| *Citrobacter koseri (diversus)* | April | ASB | Ck 19 |
| *Citrobacter koseri (diversus)* | April | ASB | Ck 20 |
| *Pseudomonas aeruginosa* | April | ASB | Pa 26 |
| *Citrobacter freundii* | April | ASB | Cf 4 |
| *Staphylococcus aureus* | April | ASB | Sa 9 |

| | | | |
|---|---|---|---|
| *Enterococcus faecalis* | October | UTI | Ef 117 |
| *Enterococcus gallinarum/casseliflavus* | November | UTI | Egc 1 |
| *Enterococcus faecalis* | November | UTI | Ef 118 |
| *Enterococcus avium* | November | UTI | Ea 3 |
| *Enterococcus faecalis* | November | UTI | Ef 129 |
| *Enterococcus faecalis* | November | UTI | Ef 130 |
| *Enterococcus faecalis* | December | UTI | Ef 131 |
| *Enterococcus faecalis* | December | UTI | Ef 132 |
| *Enterococcus faecalis* | December | UTI | Ef 133 |
| *Enterococcus faecalis* | December | UTI | Ef 134 |
| *Enterococcus faecalis* | November | UTI | Ef 128 |

| | | | |
|---|---|---|---|
| *Providencia rettgeri* | April | ASB | Pr 2 |
| *Kerstersia gyiorum* | April | ASB | Kg 1 |
| *Proteus vulgaris/penneri* | April | ASB | Pvp 7 |
| *Proteus vulgaris/penneri* | April | ASB | Pvp 8 |
| *Pseudomonas aeruginosa* | April | ASB | Pa 27 |
| *Morganella morganii* | April | ASB | Mm 11 |
| *Proteus mirabilis* | April | ASB | Pm 12 |
| *Pseudomonas aeruginosa* | April | ASB | n/a |
| *Citrobacter koseri (diversus)* | April | UTI | Ck 22 |
| *Pseudomonas aeruginosa* | April | UTI | Pa 32 |
| *Morganella morganii* | April | UTI | Mm 12 |
| *Pseudomonas aeruginosa* | April | UTI | Pa 33 |
| *Citrobacter koseri (diversus)* | April | UTI | Ck 23 |
| *Morganella morganii* | April | UTI | Mm 13 |
| *Enterobacter cloacae* | April | UTI | Enc 3.2 |
| *Enterobacter cloacae* | May | UTI | Enc 1.2 |
| *Citrobacter koseri (diversus)* | May | UTI | Ck 24 |
| *Enterobacter cloacae* | May | UTI | Enc 2.2 |
| *citrobacter koseri (diversus)* | May | UTI | Ck 25 |
| *corynebacterium jeikeium* | May | UTI | Cj 1 |
| *enterobacter cloacae* | May | UTI | Enc 5 |
| *enterobacter cloacae* | May | UTI | Enc 4 |
| *Pseudomonas aeruginosa* | May | UTI | Pa 34 |
| *Proteus vulgaris* | May | UTI | Pvp 10 |
| *Pseudomonas aeruginosa* | June | UTI | Pa 35 |
| *Enterobacter cloacae* | June | UTI | Enc 6 |
| *Pseudomonas aeruginosa* | July | UTI | Pa 36 |
| *Serratia marcescens* | July | UTI | Sms 8 |
| *Staphylococcus aureus* | July | UTI | Sa 11 |
| *Proteus mirabilis* | July | UTI | Pm 14 |
| *Enterobacter cloacae* | July | UTI | Enc 12 |
| *Enterobacter cloacae* | July | UTI | Enc 8 |
| *Enterobacter cloacae* | July | UTI | Enc 13 |
| *Proteus mirabilis* | August | UTI | Pm 16 |
| *Proteus vulgaris* | August | UTI | Pvp 11 |
| *Proteus mirabilis* | August | UTI | Pm 15 |
| *Pseudomonas aeruginosa* | August | UTI | Pa 37 |
| *Citrobacter freundii* | August | UTI | Cf 5 |
| *Pseudomonas aeruginosa* | August | UTI | n/a |
| *Pseudomonas aeruginosa* | August | UTI | Pa 39 |
| *Proteus mirabilis* | August | UTI | Pm 17 |
| *Pseudomonas aeruginosa* | August | UTI | Pa 41 |
| *Pseudomonas aeruginosa* | August | UTI | Pa 43 |
| *Acinetobacter ursingii* | August | UTI | Au 1 |
| *Stenotrophomonas maltophilia* | August | UTI | Sm 3 |
| *Citrobacter koseri (diversus)* | August | UTI | Ck 27 |
| *Citrobacter freundii* | August | UTI | Cf 6 |
| *Pseudomonas aeruginosa* | August | UTI | Pa 42 |
| *Pseudomonas aeruginosa* | August | UTI | Pa 44 |
| *Enterobacter cloacae* | August | UTI | Enc 14 |
| *Pseudomonas aeruginosa* | August | UTI | Pa 40 |
| *Proteus mirabilis* | August | UTI | Pm 18 |
| *Staphylococcus aureus* | August | UTI | Sa 12 |
| *Enterobacter cloacae* | August | UTI | Enc 15 |
| *Acinetobacter baumannii* | August | UTI | Ab 5 |
| *Serratia marcescens* | August | UTI | Sms 9.1 |
| *Pseudomonas aeruginosa* | August | UTI | Pa 45 |
| *Pseudomonas aeruginosa* | Sepetember | UTI | Pa 46 |
| *Acinetobacter baumannii* | Sepetember | UTI | Ab 6 |
| *Pseudomonas aeruginosa* | Sepetember | UTI | Pa 48 |
| *Actinotignum (Actinobaculum) schaalii* | October | UTI | As 4 |
| *Citrobacter freundii* | October | UTI | Cf 7.1 |
| *Pseudomonas aeruginosa* | October | UTI | Pa 47 |
| *Pseudomonas aeruginosa* | October | UTI | Pa 49 |
| *Pseudomonas aeruginosa* | October | UTI | Pa 50 |
| *Citrobacter freundii* | October | UTI | Cf 8 |
| *Pseudomonas aeruginosa* | October | UTI | Pa 51 |
| *Citrobacter freundii* | October | UTI | Cf 9 |
| *Pseudomonas aeruginosa* | October | UTI | Pa 52 |
| *Enterobacter cloacae* | October | UTI | Enc 18 |
| *Proteus mirabilis* | October | UTI | Pm 19 |
| *Morganella morganii* | October | UTI | Mm 14 |
| *Pseudomonas aeruginosa* | November | UTI | Pa 53 |
| *Enterobacter cloacae* | November | UTI | Enc 19 |
| *Enterobacter cloacae* | November | UTI | Enc 20 |
| *Staphylococcus aureus* | November | UTI | Sa 13 |
| *Morganella morganii* | November | UTI | Mm 15 |
| *Actinotignum (Actinobaculum) schaalii* | November | UTI | As 5 |
| *Pseudomonas aeruginosa* | November | UTI | Pa 54 |
| *Proteus vulgaris* | November | UTI | Pvp 12 |
| *Proteus vulgaris* | December | UTI | Pv 2 |
| *Providencia rettgeri* | December | UTI | Pr 3 |
| *Kerstersia gyiorum* | December | UTI | Kg 2 |
| *Serratia marcescens* | December | UTI | Sms 9.2 |
| *Citrobacter farmeri* | December | UTI | Cfa1 |
| *Citrobacter freundii* | December | UTI | Cf 7.2 |
| *Enterobacter cloacae* | August | UTI | Enc 16 |

**Table S1. The Zurich Uropathogen Collection.** A total of 663bacterial stains were identified and cultured from urine specimens of patients from the Department of Neuro-Urology, Balgrist University Hospital, Zürich, acquired between January to December 2020 and provided after routine testing by the team of the Institute of Medical Microbiology (IMM), University of Zürich. Isolates of (**a**) *E. coli*, (**b**) *Klebsiella* spp. and (**c**) *Enterococcus* spp. are shown in separate sub-tables. All other isolated species are summarized in (**d**). An overview of the species distribution can be found in **Fig. S1**. ASB = asymptomatic bacteriuria.

**Table S2. Strains used in this study.** Sources: 1 = Zurich Uropathogen Collection; 2 = in house strain collection; 3= Collection of the Institute of Veterinary Bacteriology, University of Bern, Switzerland, 4= Wissing et. al.; 5= The National Reference Center for Emerging Antibiotic Resistance (NARA), University of Fribourg, Switzerland. 6= National Reference Centre for Enteropathogenic Bacteria and Listeria (NENT) (Zurich, Switzerland)

**Escherichia coli**

| Bacterial species | Source | Designation |
|---|---|---|
| *Escherichia coli* | 1 | Ec3 |
| *Escherichia coli* | 1 | Ec14 |
| *Escherichia coli* | 1 | Ec16 |
| *Escherichia coli* | 1 | Ec20 |
| *Escherichia coli* | 1 | Ec21 |
| *Escherichia coli* | 1 | Ec22 |
| *Escherichia coli* | 1 | Ec28 |
| *Escherichia coli* | 1 | Ec32 |
| *Escherichia coli* | 1 | Ec33 |
| *Escherichia coli* | 1 | Ec34 |
| *Escherichia coli* | 1 | Ec41 |
| *Escherichia coli* | 1 | Ec42 |
| *Escherichia coli* | 1 | Ec43 |
| *Escherichia coli* | 1 | Ec46 |
| *Escherichia coli* | 1 | Ec47 |
| *Escherichia coli* | 1 | Ec52 |
| *Escherichia coli* | 1 | Ec54 |
| *Escherichia coli* | 1 | Ec56 |
| *Escherichia coli* | 1 | Ec57 |
| *Escherichia coli* | 1 | Ec62 |
| *Escherichia coli* | 1 | Ec63 |
| *Escherichia coli* | 1 | Ec65 |
| *Escherichia coli* | 1 | Ec70 |
| *Escherichia coli* | 1 | Ec74 |
| *Escherichia coli* | 1 | Ec76 |
| *Escherichia coli* | 1 | Ec77 |
| *Escherichia coli* | 1 | Ec79 |
| *Escherichia coli* | 1 | Ec82 |
| *Escherichia coli* | 1 | Ec92 |
| *Escherichia coli* | 1 | Ec93 |
| *Escherichia coli* | 1 | Ec96 |
| *Escherichia coli* | 1 | Ec97 |
| *Escherichia coli* | 1 | Ec101 |
| *Escherichia coli* | 1 | Ec117 |
| *Escherichia coli* | 1 | Ec131 |
| *Escherichia coli* | 1 | Ec132 |
| *Escherichia coli* | 1 | Ec136 |
| *Escherichia coli* | 1 | Ec137 |
| *Escherichia coli* | 1 | Ec138 |
| *Escherichia coli* | 1 | Ec139 |
| *Escherichia coli* | 1 | Ec140 |
| *Escherichia coli* | 1 | Ec141 |
| *Escherichia coli* | 1 | Ec142 |
| *Escherichia coli* | 1 | Ec143 |
| *Escherichia coli* | 1 | Ec145 |
| *Escherichia coli* | 1 | Ec146 |
| *Escherichia coli* | 1 | Ec147 |
| *Escherichia coli* | 1 | Ec148 |
| *Escherichia coli* | 1 | Ec149 |
| *Escherichia coli* | 1 | Ec151 |
| *Escherichia coli* | 1 | Ec158 |
| *Escherichia coli* | 2 | BL21 |

***Klebsiella* spp.**

| Bacterial species | Source | Designation |
|---|---|---|
| *Klebsiella aerogenes* | 1 | k92 |
| *Klebsiella aerogenes* | 1 | k112 |
| *Klebsiella oxytoca* | 1 | Ko38 |
| *Klebsiella oxytoca* | 1 | Ko68 |
| *Klebsiella oxytoca* | 1 | Ko87 |
| *Klebsiella oxytoca* | 1 | Ko101 |
| *Klebsiella pneumoniae* | 1 | KpGe (Kp1) |
| *Klebsiella pneumoniae* | 1 | Kp3 |
| *Klebsiella pneumoniae* | 1 | Kp5 |
| *Klebsiella pneumoniae* | 1 | Kp6 |
| *Klebsiella pneumoniae* | 1 | Kp8 |
| *Klebsiella pneumoniae* | 1 | Kp11 |
| *Klebsiella pneumoniae* | 1 | Kp13 |
| *Klebsiella pneumoniae* | 1 | Kp14 |
| *Klebsiella pneumoniae* | 1 | Kp15 |
| *Klebsiella pneumoniae* | 1 | Kp16 |
| *Klebsiella pneumoniae* | 1 | Kp21 |
| *Klebsiella pneumoniae* | 1 | Kp22 |
| *Klebsiella pneumoniae* | 1 | Kp26 |
| *Klebsiella pneumoniae* | 1 | Kp31 |
| *Klebsiella pneumoniae* | 1 | Kp33 |
| *Klebsiella pneumoniae* | 1 | Kp34 |
| *Klebsiella pneumoniae* | 1 | Kp36 |
| *Klebsiella pneumoniae* | 1 | Kp37 |
| *Klebsiella pneumoniae* | 1 | Kp39 |
| *Klebsiella pneumoniae* | 1 | Kp43 |
| *Klebsiella pneumoniae* | 1 | Kp45 |
| *Klebsiella pneumoniae* | 1 | Kp48 |
| *Klebsiella pneumoniae* | 1 | Kp51 |
| *Klebsiella pneumoniae* | 1 | Kp59 |
| *Klebsiella pneumoniae* | 1 | Kp66 |
| *Klebsiella pneumoniae* | 1 | Kp67 |
| *Klebsiella pneumoniae* | 1 | Kp69 |
| *Klebsiella pneumoniae* | 1 | Kp70 |
| *Klebsiella pneumoniae* | 1 | Kp71 |
| *Klebsiella pneumoniae* | 1 | Kp72 |
| *Klebsiella pneumoniae* | 1 | Kp76 |
| *Klebsiella pneumoniae* | 1 | Kp78 |
| *Klebsiella pneumoniae* | 1 | Kp79 |
| *Klebsiella pneumoniae* | 1 | Kp80 |
| *Klebsiella pneumoniae* | 1 | Kp81 |
| *Klebsiella pneumoniae* | 1 | Kp82 |
| *Klebsiella pneumoniae* | 1 | Kp83 |
| *Klebsiella pneumoniae* | 1 | Kp88 |
| *Klebsiella pneumoniae* | 1 | Kp89 |
| *Klebsiella pneumoniae* | 1 | Kp90 |
| *Klebsiella pneumoniae* | 1 | Kp93 |
| *Klebsiella pneumoniae* | 1 | Kp94 |
| *Klebsiella pneumoniae* | 1 | Kp95 |
| *Klebsiella pneumoniae* | 1 | Kp96 |
| *Klebsiella pneumoniae* | 1 | Kp97 |
| *Klebsiella pneumoniae* | 1 | Kp98 |

***Enterococcus* spp.**

| Bacterial species | Source | Designation |
|---|---|---|
| *Enterococcus faecalis* | 1 | Ef2 |
| *Enterococcus faecalis* | 1 | Ef3 |
| *Enterococcus faecalis* | 1 | Ef4 |
| *Enterococcus faecalis* | 1 | Ef5 |
| *Enterococcus faecalis* | 1 | Ef7 |
| *Enterococcus faecalis* | 1 | Ef8 |
| *Enterococcus faecalis* | 1 | Ef12 |
| *Enterococcus faecalis* | 1 | Ef14 |
| *Enterococcus faecalis* | 1 | Ef15 |
| *Enterococcus faecalis* | 1 | Ef18 |
| *Enterococcus faecalis* | 1 | Ef20 |
| *Enterococcus faecalis* | 1 | Ef24 |
| *Enterococcus faecalis* | 1 | Ef25 |
| *Enterococcus faecalis* | 1 | Ef26 |
| *Enterococcus faecalis* | 1 | Ef27 |
| *Enterococcus faecalis* | 1 | Ef28 |
| *Enterococcus faecalis* | 1 | Ef30 |
| *Enterococcus faecalis* | 1 | Ef33 |
| *Enterococcus faecalis* | 1 | Ef34 |
| *Enterococcus faecalis* | 1 | Ef36 |
| *Enterococcus faecalis* | 1 | Ef38 |
| *Enterococcus faecalis* | 1 | Ef44 |
| *Enterococcus faecalis* | 1 | Ef45 |
| *Enterococcus faecalis* | 1 | Ef46 |
| *Enterococcus faecalis* | 1 | Ef47 |
| *Enterococcus faecalis* | 1 | Ef48 |
| *Enterococcus faecalis* | 1 | Ef49 |
| *Enterococcus faecalis* | 1 | Ef51 |
| *Enterococcus faecalis* | 1 | Ef54 |
| *Enterococcus faecalis* | 1 | Ef55 |
| *Enterococcus faecalis* | 1 | Ef57 |
| *Enterococcus faecalis* | 1 | Ef58 |
| *Enterococcus faecalis* | 1 | Ef59 |
| *Enterococcus faecalis* | 1 | Ef60 |
| *Enterococcus faecalis* | 1 | Ef66 |
| *Enterococcus faecalis* | 1 | Ef67 |
| *Enterococcus faecalis* | 1 | Ef69 |
| *Enterococcus faecalis* | 1 | Ef70 |
| *Enterococcus faecalis* | 1 | Ef71 |
| *Enterococcus faecalis* | 1 | Ef72 |
| *Enterococcus faecalis* | 1 | Ef73 |
| *Enterococcus faecalis* | 1 | Ef74 |
| *Enterococcus faecalis* | 1 | Ef78 |
| *Enterococcus faecalis* | 1 | Ef79 |
| *Enterococcus faecalis* | 1 | Ef81 |
| *Enterococcus faecalis* | 1 | Ef82 |
| *Enterococcus faecalis* | 1 | Ef83 |
| *Enterococcus faecalis* | 1 | Ef88 |
| *Enterococcus faecalis* | 1 | Ef89 |
| *Enterococcus faecalis* | 1 | Ef90 |
| *Enterococcus faecalis* | 1 | Ef92 |
| *Enterococcus faecalis* | 1 | Ef93 |
| *Enterococcus faecalis* | 1 | Ef94 |
| *Enterococcus faecalis* | 1 | Ef95 |
| *Enterococcus faecalis* | 1 | Ef96 |
| *Enterococcus faecalis* | 2 | JH2-2 |
| *Enterococcus faecium* | 5 | NARA388 |
| *Enterococcus faecium* | 3 | LPK0102 |
| *Enterococcus faecium* | 3 | DVT02299 |
| *Enterococcus faecium* | 3 | LPK0229 |
| *Enterococcus faecium* | 3 | LPK1232 |
| *Enterococcus faecium* | 3 | LPK3080 |
| *Enterococcus faecium* | 3 | LPK9233 |
| *Enterococcus faecium* | 4 | AW-51B |
| *Enterococcus faecium* | 4 | AW-54B |
| *Enterococcus faecium* | 4 | AW-51D |
| *Enterococcus faecium* | 4 | AW-17A |
| *Enterococcus faecium* | 4 | AW-22E |
| *Enterococcus faecium* | 4 | AW-32A |

**Other Gram-positives**

| Bacterial species | Source | Designation |
|---|---|---|
| *Corynebacterium jeikeium* | 1 | Cj1 |
| *Lactobacillus acidophilus* | 2 | ATCC4356 |
| *Lactobacillus acidophilus* | 2 | DSM20079T |
| *Lactobacillus casei* | 2 | ATCC334 |
| *Lactococcus lactis* | 2 | Bu2 |
| *Lactococcus lactis* | 2 | K214 |
| *Micrococcus kristinae* | 2 | DSM20032 |
| *Pediococcus acidilactici* | 2 | DSM20284 |
| *Staphylococcus aureus* | 2 | Newman |
| *Staphylococcus aureus* | 1 | Sa2 |
| *Staphylococcus aureus* | 1 | Sa5 |
| *Staphylococcus aureus* | 1 | Sa6 |
| *Staphylococcus aureus* | 1 | Sa7 |
| *Staphylococcus aureus* | 1 | Sa8 |
| *Staphylococcus aureus* | 1 | Sa9 |
| *Staphylococcus aureus* | 1 | Sa10 |
| *Staphylococcus aureus* | 1 | Sa11 |
| *Staphylococcus aureus* | 1 | Sa12 |
| *Staphylococcus aureus* | 1 | Sa13 |
| *Staphylococcus aureus* | 2 | PSK |
| *Staphylococcus epidermidis* | 2 | S602 |
| *Staphylococcus epidermidis* | 2 | S414 |
| *Streptococcus mutans* | 2 | OMZ381 |
| *Streptococcus mutans* | 2 | OMZ7 |
| *Streptococcus mutans* | 2 | OMZ25 |
| *Streptococcus mutans* | 2 | OMZ27 |
| *Streptococcus mutans* | 2 | OMZ30 |
| *Streptococcus pneumoniae* | 2 | R6 |
| *Streptococcus pyogenes* | 2 | ATCC700294 |
| *Streptococcus salivarius* | 2 | DSM20560T |

**Other Gram-negatives**

| Bacterial species | Source | Designation |
|---|---|---|
| *Acinetobacter baumanii* | 1 | Ab4 |
| *Acinetobacter baumanii* | 1 | Ab6 |
| *Citrobacter farmeri* | 1 | Cfa1 |
| *Citrobacter freundii* | 1 | Cf5 |
| *Citrobacter koseri* | 1 | Ck1 |
| *Citrobacter koseri* | 1 | Ck21 |
| *Citrobacter koseri* | 1 | Ck25 |
| *Enterobacter cloacae* | 1 | Ecl3 |
| *Enterobacter cloacae* | 1 | Ecl10 |
| *Enterobacter cloacae* | 1 | Ecl6 |
| *Enterobacter cloacae* | 1 | Enc6 |
| *Kerstersia gyiorum* | 1 | Kg2 |
| *Morganella morganii* | 1 | Mm8 |
| *Morganella morganii* | 1 | Mm13 |
| *Proteus mirabilis* | 1 | Pm13 |
| *Proteus mirabilis* | 1 | Pm5 |
| *Proteus vulgaris* | 1 | Pv3 |
| *Providencia rettgeri* | 1 | Pr3 |
| *Pseudomonas aeruginosa* | 1 | Pa38 |
| *Pseudomonas aeruginosa* | 1 | Pa28 |
| *Pseudomonas aeruginosa* | 1 | Pa34 |
| *Raoutella ornithinolytica* | 1 | Ro1 |
| *Raoutella ornithinolytica* | 1 | Ro2 |
| *Salmonella* Enteritidis | 6 | N58-09 |
| *Salmonella* Infantis | 6 | N63-09 |
| *Salmonella* Newport | 6 | N2821-08 |
| *Salmonella* Senftenberg | 6 | N1918-08 |
| *Salmonella* Typhimurium | 2 | DB 7155 |
| *Serratia marcescens* | 1 | Sm5 |
| *Stenotrophomonas maltophilia* | 1 | Sma3 |

**Table S3: Bacteria isolated during the field evaluation from patient urine**. The source of the urine corresponds to the patient number. Nr 1-147: Site 1 (Balgrist University Hospital, Zurich, Switzerland); Nr. 148-206, Site 2 (University Hospital Zurich, Zurich, Switzerland).

| ***Escherichia coli*** | | | ***Klebsiella* spp.** | | | ***Enterococcus* spp.** | | |
|---|---|---|---|---|---|---|---|---|
| **Species designation** | **Strain designation** | **Patient Nr. field evaluation** | **Species designation** | **Strain designation** | **Patient Nr. field evaluation** | **Species designation** | **Strain designation** | **Patient Nr. field evaluation** |
| *Eschericia coli* | BAL Ec1 | 6 | *Klebsiella aerogenes* | USZ Ka11 | 152 | *Enterococcus avium* | USZ Ea08 | 155 |
| *Eschericia coli* | BAL Ec2 | 7 | *Klebsiella aerogenes* | USZ Ka53 | 192 | *Enterococcus faecalis* | BAL Ef1 | 1 |
| *Eschericia coli* | BAL Ec3 | 8 | *Klebsiella oxytoca* | BAL Ko1 | 27 | *Enterococcus faecalis* | BAL Ef2 | 7 |
| *Eschericia coli* | BAL Ec4 | 13 | *Klebsiella oxytoca* | BAL Ko2 | 48 | *Enterococcus faecalis* | BAL Ef3 | 36 |
| *Eschericia coli* | BAL Ec5 | 14 | *Klebsiella oxytoca* | BAL Ko3 | 65 | *Enterococcus faecalis* | BAL Ef4 | 41 |
| *Eschericia coli* | BAL Ec6 | 17 | *Klebsiella oxytoca* | BAL Ko4 | 138 | *Enterococcus faecalis* | BAL Ef5 | 43 |
| *Eschericia coli* | BAL Ec7 | 19 | *Klebsiella oxytoca* | USZ Ko09 | 150 | *Enterococcus faecalis* | BAL Ef6 | 57 |
| *Eschericia coli* | BAL Ec8 | 23 | *Klebsiella oxytoca* | USZ Ko51 | 190 | *Enterococcus faecalis* | BAL Ef7 | 63 |
| *Eschericia coli* | BAL Ec9 | 24 | *Klebsiella pneumoniae* | BAL Kp1 | 11 | *Enterococcus faecalis* | BAL Ef8 | 76 |
| *Eschericia coli* | BAL Ec10 | 26 | *Klebsiella pneumoniae* | BAL Kp2 | 26 | *Enterococcus faecalis* | BAL Ef9 | 89 |
| *Eschericia coli* | BAL Ec11 | 35 | *Klebsiella pneumoniae* | BAL Kp3 | 33 | *Enterococcus faecalis* | BAL Ef10 | 96 |
| *Eschericia coli* | BAL Ec12 | 41 | *Klebsiella pneumoniae* | BAL Kp4 | 63 | *Enterococcus faecalis* | BAL Ef19 | 107 |
| *Eschericia coli* | BAL Ec13 | 43 | *Klebsiella pneumoniae* | BAL Kp5 | 76 | *Enterococcus faecalis* | BAL Ef11 | 110 |
| *Eschericia coli* | BAL Ec14 | 45 | *Klebsiella pneumoniae* | BAL Kp6 | 77 | *Enterococcus faecalis* | BAL Ef12 | 113 |
| *Eschericia coli* | BAL Ec15 | 48 | *Klebsiella pneumoniae* | BAL Kp7 | 84 | *Enterococcus faecalis* | BAL Ef13 | 119 |
| *Eschericia coli* | BAL Ec16 | 53 | *Klebsiella pneumoniae* | BAL Kp8 | 87 | *Enterococcus faecalis* | BAL Ef14 | 126 |
| *Eschericia coli* | BAL Ec17 | 56 | *Klebsiella pneumoniae* | BAL Kp9 | 116 | *Enterococcus faecalis* | BAL Ef15 | 129 |
| *Eschericia coli* | BAL Ec18 | 57 | *Klebsiella pneumoniae* | BAL Kp10 | 117 | *Enterococcus faecalis* | BAL Ef16 | 131 |
| *Eschericia coli* | BAL Ec19 | 64 | *Klebsiella pneumoniae* | BAL Kp11 | 141 | *Enterococcus faecalis* | BAL Ef17 | 139 |
| *Eschericia coli* | BAL Ec20 | 75 | *Klebsiella pneumoniae* | BAL Kp12 | 144 | *Enterococcus faecalis* | BAL Ef18 | 144 |
| *Eschericia coli* | BAL Ec21 | 78 | *Klebsiella pneumoniae* | USZ Kp08 | 149 | *Enterococcus faecalis* | USZ Ef08 | 155 |
| *Eschericia coli* | BAL Ec22 | 79 | *Klebsiella pneumoniae* | USZ Kp37 | 176 | *Enterococcus faecalis* | USZ Ef09 | 156 |
| *Eschericia coli* | BAL Ec23 | 80 | *Klebsiella pneumoniae* | USZ Kp38 | 177 | *Enterococcus faecalis* | USZ Ef14 | 161 |
| *Eschericia coli* | BAL Ec24 | 86 | *Klebsiella pneumoniae* | USZ Kp41 | 180 | *Enterococcus faecalis* | USZ Ef15 | 162 |
| *Eschericia coli* | BAL Ec25 | 88 | | | | *Enterococcus faecalis* | USZ Ef16 | 163 |
| *Eschericia coli* | BAL Ec26 | 93 | | | | *Enterococcus faecalis* | USZ Ef28 | 175 |
| *Eschericia coli* | BAL Ec27 | 108 | | | | *Enterococcus faecalis* | USZ Ef32 | 179 |
| *Eschericia coli* | BAL Ec28 | 111 | | | | *Enterococcus faecalis* | USZ Ef33 | 180 |
| *Eschericia coli* | BAL Ec29 | 114 | | | | *Enterococcus faecalis* | USZ Ef38 | 185 |
| *Eschericia coli* | BAL Ec30 | 118 | | | | *Enterococcus faecalis* | USZ Ef39 | 186 |
| *Eschericia coli* | BAL Ec31 | 127 | | | | *Enterococcus faecalis* | USZ Ef40 | 187 |
| *Eschericia coli* | BAL Ec32 | 132 | | | | *Enterococcus faecalis* | USZ Ef41 | 188 |
| *Eschericia coli* | BAL Ec33 | 133 | | | | *Enterococcus faecalis* | USZ Ef43 | 190 |
| *Eschericia coli* | BAL Ec34 | 134 | | | | *Enterococcus faecalis* | USZ Ef46 | 193 |
| *Eschericia coli* | BAL Ec35 | 139 | | | | *Enterococcus faecalis* | USZ Ef48 | 195 |
| *Eschericia coli* | USZ Ec01 | 146 | | | | *Enterococcus faecalis* | USZ Ef56 | 203 |
| *Eschericia coli* | USZ Ec03 | 147 | | | | *Enterococcus faecalis* | USZ Ef59 | 206 |
| *Eschericia coli* | USZ Ec07 | 148 | | | | | | |
| *Eschericia coli* | USZ Ec12 | 153 | | | | | | |
| *Eschericia coli* | USZ Ec14 | 155 | | | | | | |
| *Eschericia coli* | USZ Ec24 | 165 | | | | | | |
| *Eschericia coli* | USZ Ec33 | 172 | | | | | | |
| *Eschericia coli* | USZ Ec38 | 177 | | | | | | |
| *Eschericia coli* | USZ Ec39 | 178 | | | | | | |
| *Eschericia coli* | USZ Ec40 | 179 | | | | | | |
| *Eschericia coli* | USZ Ec41 | 180 | | | | | | |
| *Eschericia coli* | USZ Ec42 | 181 | | | | | | |
| *Eschericia coli* | USZ Ec53 | 192 | | | | | | |
| *Eschericia coli* | USZ Ec56 | 195 | | | | | | |
| *Eschericia coli* | USZ Ec59 | 198 | | | | | | |

**Table S4: Plasmids used and constructed during this study**. P1 = Primer 1, P2 = Primer 2, *cps* = major capsid protein gene, *nluc* = nanoluciferase gene.

| Name | Fragment | Template | P1 | P2 | Reference | Application |
|---|---|---|---|---|---|---|
| pLEB579 | -<br>- | -<br>- | -<br>- | -<br>- | Beasley et al., 2004 | backbone Gram-positives |
| pCas9 | -<br>- | -<br>- | -<br>- | -<br>- | Jiang et al., 2013 | backbone Gram-negatives |
| pUC19 | -<br>- | -<br>- | -<br>- | -<br>- | Yanisch-Perron et al., 1985 | backbone Gram-negatives |
| pRSF-deut1 | -<br>- | -<br>- | -<br>- | -<br>- | Novagen, Merck | backbone Gram-negatives |
| pUC19-$Kan^R$ | F1<br>F2 | pRSF-deut1<br>pUC19 | P37<br>P39 | P38<br>P40 | this study | backbone Gram-negatives |
| $pCas9_{T4}$ | F1<br>F2 | pCas9<br>$String_{T4-RSRSR}$ | P5<br>P7 | P6<br>P8 | this study | spacer pair targeting *cps* of phage T4, scrambled (scr) control vector in Gram-negatives |
| $pSelect_E$ | F1<br>F2 | pCas9<br>$String_{E-RSRSR}$ | P5<br>P7 | P6<br>P8 | this study | spacer pair targeting *cps* and *gp170* of phage E2/E4 |
| $pSelect_K$ | F1<br>F2 | pCas9<br>$String_{K-RSRSR}$ | P5<br>P7 | P6<br>P8 | this study | spacer pair targeting *gp167* and *gp168* of phage K1 (= spacer pair targeting *gp168* and *gp169* of phage K4) |
| $pSelect_{T4}$ | F1<br>F2 | pLEB579<br>$pCas9_{T4}$ | P1<br>P3 | P2<br>P4 | this study | spacer pair targeting *cps* of phage T4, scrambled (scr) control vector in Gram-positives |
| $pSelect_{EfS3}$ | F1<br>F2 | $pSelect_{T4}$<br>$String_{EfS3-RSRSR}$ | P5<br>P7 | P6<br>P8 | this study | spacer pair targeting *cps* and of phage EfS3 |
| $pSelect_{EfS7}$ | F1<br>F2 | $pSelect_{T4}$<br>$String_{EfS7-RSRSR}$ | P5<br>P7 | P6<br>P8 | this study | spacer pair targeting *cps* of phage EfS7 |
| $pEdit_{EfS3}$ | F1<br>F2 | pLEB579<br>$String_{EfS3-nluc}$ | P9<br>P11 | P10<br>P12 | this study | Editing template for site-directed *nluc* insertion downstream of the *cps* gene of phage EfS3 |
| $pEdit_{EfS7}$ | F1<br>F2 | pLEB579<br>$String_{EfS7-nluc}$ | P13<br>P15 | P14<br>P16 | this study | Editing template for site-directed *nluc* insertion downstream of the *cps* gene of phage EfS7 |
| $pEdit_E$ | F1<br>F2 | pUC19<br>$String_{E-nluc}$ | P29<br>P31 | P30<br>P32 | this study | Editing template for site-directed nluc insertion downstream of the *cps* gene of phage E2/E4 |
| $pEdit_K$ | F1<br>F2 | pUC19-$Kan^R$<br>$String_{K-nluc}$ | P33<br>P35 | P34<br>P36 | this study | Editing template for site-directed *nluc* insertion downstream of the prohead assembly protein gene of phage K1/K4 |

**Table S5: Primer and synthetic DNA string sequences used in this study.** Underlined = repeat sequence; red = spacer sequence; red+bold+underlined = PAM mutation; bold+underlined = protospacer sequence; bold italics = ribosomal binding site; blue+bold = codon-optimized *nluc* sequence.

| Nucleotide name | Sequence 5'-3' | Application |
|---|---|---|
| P1 | GACAGCATCGCCAGTCACTATTCTAGACTCGAGTCGATGTTAAAC | pSelect, cloning |
| P2 | CTGACTGCGTTAGCAATTTGATATCGCGCGCTATTAATCGCAAC | pSelect, cloning |
| P3 | AAATTGCTAACGCAGTCAGG | pSelect, cloning |
| P4 | ATAGTGACTGGCGATGCT | pSelect, cloning |
| P5 | TCAGCTAGACTTCAGTCTTGAAAAG | pSelect, cloning |
| P6 | GACTCCATTCAACATTGCCGA | pSelect, cloning |
| P7 | GCAGTAATACAGGGGCTTTTC | pSelect, cloning |
| P8 | CCTCTTTCTCAAGTTATCATCGG | pSelect, cloning |
| P9 | AGAAAAACCAAAAGCAGTCGATGTTAAACCGTGTGCTCTACG | pEdit$_{EfS3}$, cloning |
| P10 | GATTTGGTAGAACAACGCGCTATTAATCGCAACATCAAACC | pEdit$_{EfS3}$, cloning |
| P11 | GCGATTAATAGCGCGTTGTTCTACCAAATCGCACG | pEdit$_{EfS3}$, cloning |
| P12 | GGTTTAACATCGACTGCTTTTGGTTTTTCTTCTTTTAC | pEdit$_{EfS3}$, cloning |
| P13 | AGTCGATGTTAAACCGTGTGCTCTACG | pEdit$_{EfS7}$, cloning |
| P14 | CGCGCTATTAATCGCAACATCAAACC | pEdit$_{EfS7}$, cloning |
| P15 | GGGGCTTTTATTTTGGTTTGATGTTG | pEdit$_{EfS7}$, cloning |
| P16 | TTATAGTTTTGGTCGTAGAGCACACG | pEdit$_{EfS7}$, cloning |
| P17 | GAGAAATGGAAGTTGAATTAAG | pSelect$_{EfS}$, pEdit$_{EfS}$, verification |
| P18 | GATAATGAACTGTGCTGATTAC | pSelect$_{EfS}$, pEdit$_{EfS}$, verification |
| P19 | ACGATGCAGAATCTGCTCC | EfS3, verification |
| P20 | GGAACAGTTAGACCGTTCC | EfS3, verification |
| P21 | AAATGACTGATAGCTACGAGTG | EfS7, verfification |
| P22 | TTTGCTTCCGTCAGAAGC | EfS7, verfification |
| P23 | GGTCTGTATTTCCCAGAACC | pSelect, verification |
| P24 | AAGGTTCTGGGAAATACAGACCG | pSelect, verification |
| P25 | TAAATACTGAAATAACTAAGGCTCCC | pSelect, verification |
| P26 | AGATAGATTTAATGCTTCATTAGGTACC | pSelect, verification |
| P27 | ATTCAATAGACAATAAGGTCTTAACGC | pSelect, verification |
| P28 | AAATTCGGACAAGCTTATTGCTCG | pSelect, verification |
| P29 | TGCAGCATAAGAAATGCCGGGCCTGGGGTGCCTAATGA | pEdit$_{E}$, cloning |
| P30 | TTAACCCGGATAACGAATTTTAAGCCAGCCCCGACAC | pEdit$_{E}$, cloning |
| P31 | ACTCATTAGGCACCCCAGGCCCGGCATTTCTTATGCTG | pEdit$_{E}$, cloning |
| P32 | CGGGTGTCGGGGCTGGCTTAAAATTCGTTATCCGGGTTAAG | pEdit$_{E}$, cloning |
| P33 | GGTGGATGGTTTAATTCTCGCCTGGGGTGCCTAATGA | pEdit$_{K}$, cloning |
| P34 | GCTGCTTCTAAATACTTTGACTAAGCCAGCCCCGACAC | pEdit$_{K}$, cloning |
| P35 | CACTCATTAGGCACCCCAGGCGAGAATTAAACCATCCACCTC | pEdit$_{K}$, cloning |
| P36 | GGGTGTCGGGGCTGGCTTAGTCAAAGTATTTAGAAGCAGCT | pEdit$_{K}$, cloning |
| P37 | AATAATATTGAAAAAGGAAGAGTATGAGCCATATTCAACGGG | pUC19-KanR, cloning |
| P38 | GAGTAAACTTGGTCTGACAGTTAGAAAAACTCATCGAGCATC | pUC19-KanR, cloning |
| P39 | CGTTGAATATGGCTCATACTCTTCCTTTTTCAATATTATTGAAG | pUC19-KanR, cloning |
| P40 | TGCTCGATGAGTTTTTCTAACTGTCAGACCAAGTTTACTCA | pUC19-KanR, cloning |
| P41 | CATTAATGCAGCTGGCACGA | pEDIT$_{E/K}$, verification |
| P42 | GTCACAGCTTGTCTGTAAGCG | pEDIT$_{E/K}$, verification |
| P43 | GGTAACTTCATTATCGCTTCC | E, verification |
| P44 | TGTGATTTGTTCTCGGTCTAC | E, verification |
| P45 | GTCTTTATCCAAAACGCATTC | K1/K4 verification |
| P46 | CAAACAGTTCAGCCTTAATAC | K1, verification |
| P47 | CAAACAGGTCAGCTTTGATA | K4, verification |
| String$_{EfS3\text{-}RSRSR}$ | AACACGCATTGATTTGAGTCAGCTAGGAGGTGACTGAAGTATATTTTAGATGAAGATTATTTCTTAATAACTAAAAATATGGTATA<br>ATACTCTTAATAAATGCAGTAATACAGGGGCTTTTCAAGACTGAAGTCTAGCTGAGACAAATAGTGCGATTACGAAATTTTTTAGA<br>CAAAAATAGTCTACGAGGTTTTAGAGCTATGCTGTTTTGAATGGTCCCAAAACAATTCACCTACGAATACGTCAGCAGTTTCAGTT<br>TTAGAGCTATGCTGTTTTGAATGGTCCCAAAACTAAAATCTACAGTATTACAAAATAAAAAAGGTTTTAGAGCTATGCTGTTTTGA<br>ATGGTCCCAAAACTTCAGCACACTGAGACTTGTTGAGTTCCATGTTTTAGAGCTATGCTGTTTTGAATGGACTCCATTCAACATTG<br>CCGATGATAACTTGAGAAAGAGGGTTAATACCAGCAGTCGGA | spacer pair targeting *cps* and of phage EfS3 |
| String$_{EfS7\text{-}RSRSR}$ | AGTATATTTTAGATGAAGATTATTTCTTAATAACTAAAAATATGGTATAATACTCTTAATAAATGCAGTAATACAGGGGCTTTTCA<br>AGACTGAAGTCTAGCTGAGACAAATAGTGCGATTACGAAATTTTTTAGACAAAAATAGTCTACGAGGTTTTAGAGCTATGCTGTTT<br>TGAATGGTCCCAAAACAACGTGCAATCTTCATCAACAAGGTAGCTGGTTTTAGAGCTATGCTGTTTTGAATGGTCCCAAAACAGAC<br>TTATAATATTTACCAAGATGGAGCGAGTTTTAGAGCTATGCTGTTTTGAATGGTCCCAAAACTTCAGCACACTGAGACTTGTTGAG<br>TTCCATGTTTTAGAGCTATGCTGTTTTGAATGGACTCCATTCAACATTGCCGATGATAACTTGAGAAAGAGGGTTAATACCAGCAG<br>TCGGATACCTTCCTATTCTTTCTGTTAAAGCGTTTTCATGTTATAATAGGCAAAAGAAGAGTAGTGTGATCGTCCATTCCGACAGC<br>ATCGCCAGTCACTAT | spacer pair targeting *cps* of phage EfS7 |
| String$_{T4\text{-}RSRSR}$ | AACACGCATTGATTTGAGTCAGCTAGGAGGTGACTGAAGTATATTTTAGATGAAGATTATTTCTTAATAACTAAAAATATGGTATA<br>ATACTCTTAATAAATGCAGTAATACAGGGGCTTTTCAAGACTGAAGTCTAGCTGAGACAAATAGTGCGATTACGAAATTTTTTAGA<br>CAAAAATAGTCTACGAGGTTTTAGAGCTATGCTGTTTTGAATGGTCCCAAAACTATACACGTCTAAAGTAAGCGTTTTTACCAGTT<br>TTAGAGCTATGCTGTTTTGAATGGTCCCAAAACGTCCTAAAACTATTCGCAATTGGATAAATGGTTTTAGAGCTATGCTGTTTTGA<br>ATGGTCCCAAAACTTCAGCACACTGAGACTTGTTGAGTTCCATGTTTTAGAGCTATGCTGTTTTGAATGGACTCCATTCAACATTG<br>CCGATGATAACTTGAGAAAGAGGGTTAATACCAGCAGTCGGA | spacer pair targeting *cps* of phage T4 |
| String$_{E\text{-}RSRSR}$ | AACACGCATTGATTTGAGTCAGCTAGGAGGTGACTGAAGTATATTTTAGATGAAGATTATTTCTTAATAACTAAAAATATGGTATA<br>ATACTCTTAATAAATGCAGTAATACAGGGGCTTTTCAAGACTGAAGTCTAGCTGAGACAAATAGTGCGATTACGAAATTTTTTAGA<br>CAAAAATAGTCTACGAGGTTTTAGAGCTATGCTGTTTTGAATGGTCCCAAAACACCCATTTGCAGAATCCGCTGCTCAGGCTCGTT<br>TTAGAGCTATGCTGTTTTGAATGGTCCCAAAACTTGGGCGACCGATTGAGTTGCTATTCGTTGGTTTTAGAGCTATGCTGTTTTGA<br>ATGGTCCCAAAACTTCAGCACACTGAGACTTGTTGAGTTCCATGTTTTAGAGCTATGCTGTTTTGAATGGACTCCATTCAACATTG<br>CCGATGATAACTTGAGAAAGAGGGTTAATACCAGCAGTCGGA | spacer pair targeting *cps* and *gp170* of phage E2/E4 |
| String$_{K\text{-}CPS\text{-}RSRSR}$ | AACACGCATTGATTTGAGTCAGCTAGGAGGTGACTGAAGTATATTTTAGATGAAGATTATTTCTTAATAACTAAAAATATGGTATA<br>ATACTCTTAATAAATGCAGTAATACAGGGGCTTTTCAAGACTGAAGTCTAGCTGAGACAAATAGTGCGATTACGAAATTTTTTAGA<br>CAAAAATAGTCTACGAGGTTTTAGAGCTATGCTGTTTTGAATGGTCCCAAAACCGAGAGCCATAAAAGCATCATCAGCACTATGTT<br>TTAGAGCTATGCTGTTTTGAATGGTCCCAAAACGAGCGACGGAAGCATCGATAGTTTTAGCTTGTTTTAGAGCTATGCTGTTTTGA<br>ATGGTCCCAAAACTTCAGCACACTGAGACTTGTTGAGTTCCATGTTTTAGAGCTATGCTGTTTTGAATGGACTCCATTCAACATTG<br>CCGATGATAACTTGAGAAAGAGGGTTAATACCAGCAGTCGGA | spacer pair targeting *gp167* and *gp168* of phage K1 (= spacer pair targeting *gp168* and *gp169* of phage K4) |
| String$_{EfS3\text{-}nluc}$ | ATTCTGCAATCATTAACGCTACTGACGGTGTTAAATTAGCAATCACTATCAACAGCATGTATCAAGCTCGTCCTCAATACGTAGCT<br>ATCTACCGTAAAGGACATCAAACAGGATTGTTCTACCAAATCGCACGTGTACCTGCAAGTAAAGCAGAAAACAACGTAATCACTTT<br>CTACGATTTGAACCAACAAATTCCTGAAACTGCTGACGTATTCGTAGGTGAATTGACTCCATCTGTAGTTCACTTGTTCGAATTAC<br>TACCAATGATGCGTTTACCATTAGCTCAAATGAACGCTTCTGTAACATTCGCAGTTCTATGGTATGGTGCTTTAGCTCTTCGTGCA<br>CCTAAGAAATGGGCACGAATCAAGAACGTTAAATACATTCCATCTGGAAACGTAATTAACGAACGCTAAGAGGAGGTAAATATATA<br>TGGTTTTCACATTAGAAGATTTCGTTGGTGATTGGCGTCAAACAGCTGGTTACAACTTAGATCAAGTTTTAGAACAAGGTGGTGTT<br>TCATCATTATTCCAAAACTTAGGTGTTTCAGTTACACCAATCCAACGTATCGTTTTATCAGGTGAAAACGGTTTAAAAATCGATAT<br>CCACGTTATCATCCCATACGAAGGTTTATCAGGTGATCAAATGGGTCAAATCGAAAAAATCTTCAAAGTTGTTTACCCAGTTGATG<br>ATCACCACTTCAAAGTTATCTTACACTACGGTACATTAGTTATCGATGGTGTTACACCAAACATGATCGATTACTTCGGTCGTCCA<br>TACGAAGGTATCGCTGTTTTCGATGGTAAAAAAATCACAGTTACAGGTACATTATGGAACGGTAACAAAATCATCGATGAACGTTT<br>AATCAACCCAGATGGTTCATTATTATTCCGTGTTACAATCAACGGTGTTACAGGTTGGCGTTTATGTGAACGTATCTTAGCTTAAT<br>CTTAGTTCAACTAAGAAAAATTGAATAGAAAACGGAATAGGGAACGGTCTAACTGTTCCCTATTTTTATTATAACGAATTAAATGG<br>AGGAATACAAATGTTAAAATCTACAGTATTACAAAATAAAAAAGTTGCAACATCTTTTGGAGAAGCTGAATTTAACCATTTAGGAG<br>AAAGTCATAACTTGCCAGAGGAACATCAAAAAGCTTTAGCTGAAAAGGTACCTTATTTGACACACATTCCTGATAAGAAAGAAAAT<br>CCTAAAGAGGAAAAAGAAGTAAAAGAAGAAAAACCAAAAGCTAAAAAAGCACCTGCTAAAGCTAAAACAGCCAAAAAAGATAAAGA<br>GTAGGAGGTCATCATATGTACCCAGATTACGGTTATGGTGATGGCGGAAACCAAGTTAACCGTGAATAT | Editing template for site-directed *nluc* insertion downstream of major capsid protein (*cps*) sequence of phage EfS3 |
| String$_{EfS7\text{-}nluc}$ | AAAGTCGAAGGGGGCTTTTATTTTGGTTTGATGTTGCGATTAATAGCGCGTGAAATGACTGATAGCTACGAGTGGGTTATTGATGC<br>TCAAGTAGCATTTGGCGAAGAGTTTGATGAAGAAACTTACTTATTCATCTCACCTAAACGCCGTGCAACAATTTTGAAATCTAAAG<br>ACTTCATTCACATTCAACAAGGCGTGTCTGTAATCAAAGGCCACTTAGGTGAAATCTACGGAATGAACATTGTAGTTTCTAACAAG<br>GTTGGTGAAGACGAAGCGTTCGTATTAAAACGTGGAGCTTTATCATTAATTATGAAACGTGACTACATGGTTGAAGAAGTGCGTGA<br>AGGTATGAAACGTCAAACTAACATCACTGCTGACCAACACTACGTAGCATTCGTTAAAGATGCTAAACGTGCAATCTTCATCAACA<br>AGGTAGCTGCAGGAAAGTAGGAGGAGGTAAATATATATGGTTTTCACATTAGAAGATTTCGTTGGTGATTGGCGTCAAACAGCTGG<br>TTACAACTTAGATCAAGTTTTAGAACAAGGTGGTGTTTCATCATTATTCCAAAACTTAGGTGTTTCAGTTACACCAATCCAACGTA<br>TCGTTTTATCAGGTGAAAACGGTTTAAAAATCGATATCCACGTTATCATCCCATACGAAGGTTTATCAGGTGATCAAATGGGTCAA<br>ATCGAAAAAATCTTCAAAGTTGTTTACCCAGTTGATGATCACCACTTCAAAGTTATCTTACACTACGGTACATTAGTTATCGATGG<br>TGTTACACCAAACATGATCGATTACTTCGGTCGTCCATACGAAGGTATCGCTGTTTTCGATGGTAAAAAAATCACAGTTACAGGTA<br>CATTATGGAACGGTAACAAAATCATCGATGAACGTTTAATCAACCCAGATGGTTCATTATTATTCCGTGTTACAATCAACGGTGTT<br>ACAGGTTGGCGTTTATGTGAACGTATCTTAGCTTAACAGCCCCCAAGAATCTTCAATCTAGCGGGGTTACTGAGGACTCAGTTACT<br>CTGACTTGGGAAGACGGAGGGGCAAGCCTCTAGTAAGCTAAACGAGAGGTTGCGGAAACGTTCCGTGACCTCTTTTACTTTATCAT<br>AAATCAGGAGGAAAAGCAATGGCAAAGACTTATAATATTTACCAAGATGGAGCGAAAGTTCAGGAAGGCATTGCAGAGTTAACTAA<br>AACCATTACAGGCCTTACGCCAAACACTTCATACAAGTTTGAGGTAACAGCTGTTGAGGACGGCGTGGAGAGTGCTAAGTCCACAG<br>CGGTAACAGCCAAAACAAACCCGCGTAAAGTAGCTACAGTAACAGCTTCTCAAAAGACTATGTCTCTAGCTTCTGACGGAAGCAAA<br>GCATTGAGTCGATGTTAAACCGTGTGCTCTACGACCAAAACTATAAAACCTTTAAG | Editing template for site-directed *nluc* insertion downstream of major capsid protein (*cps)* sequence of phage EfS7 |
| String$_{E\text{-}nluc}$ | CCGGCATTTCTTATGCTGCACAAGGTCTGGCTACTGGCTTTAACACTGATACTACCAAGTCAGTATTTGCTGGTGTTCTGGGTGGT<br>AAATACCGTGTATATATCGACCAGTATGCTAAACAGGATTATTTCACCGTAGGTTATAAAGGTCCGAACGAAATGGATGCTGGTAT<br>TTACTATGCTCCATATGTAGCTCTGACTCCGCTGCGTGGTTCTGATCCGAAGAACTTCCAACCGGTAATGGGATTCAAAACTCGTT<br>ACGGTATCGGTATCAACCCATTTGCAGAATCCGCTGCTCAGGCTCCAGCTTCTCGCATCCAGAGCGGTATGCCTTCTATTCTGAAT<br>AGCCTTGGTAAAAACGCTTACTTCCGCCGTGTATATGTTAAAGGTATCTAAAGTACGAGGAGGTAAATATATATGGTTTTCACCCT<br>GGAAGACTTCGTTGGTGACTGGCGTCAGACCGCTGGTTACAACCTCGACCAGGTGCTAGAACAGGGTGGTGTTTCTTCTTTATTCC<br>AGAACCTGGGTGTAAGCGTTACCCCGATCCAGCGTATCGTTCTGTCTGGTGAAAACGGTCTGAAAATCGACATCCACGTTATCATC<br>CCGTACGAAGGTCTGTCTGGTGACCAGATGGGTCAGATCGAAAAAATCTTCAAAGTTGTTTACCCGGTTGACGACCACCACTTCAA<br>AGTTATCCTGCACTACGGTACCCTGGTTATCGACGGTGTTACCCCGAACATGATCGACTACTTCGGTCGTCCGTACGAAGGTATCG<br>CTGTTTTCGACGGTAAAAAAATCACCGTTACCGGTACCCTGTGGAACGGTAACAAAATCATCGACGAACGTCTGATCAACCCGGAC<br>GGTTCTCTGCTGTTCCGTGTTACCATCAACGGTGTTACCGGTTGGCGTCTGTGCGAACGTATCCTGGCTTAATCTTTGACGATTTA<br>AGGGACCTTCGGGTCCCTTTAGTTGTTTTTATGGTATAAATAAATCATATAAACTGAAAGGAAAGCGCAATGGCTAAAATCAACGA<br>ACTTCTGCGCGAATCAACAACAACGAATAGCAACTCAATCGGTCGCCCAAATCTCGTTGCTTTGACTCGCGCTACCACTAAATTAA<br>TATATTCTGACATTGTAGCAACGCAAAGAACTAATCAACCTGTTGCTGCTTTTTATGGTATCAAATACCTTAACCCGGATAACGAA<br>TTT | Editing template for site-directed *nluc* insertion downstream of the *cps* gene of phage E2/E4 |
| String$_{K\text{-}nluc}$ | GAGAATTAAACCATCCACCTCGTGCAAATGTTGACCCGATGCAAGCCGCAATCATTATCGAAGATATGTGGTGGAAAGGCAACGAT<br>GTTTATGGGCGTGCAAGAATCATCGAAGGAGACCACGGTCCAGGTGATAAATTAGCCGCTAACATCCGTGCTGGTTGGATTCCTGG<br>TGTATCAAGCCGTGGGCTCGGTTCTTTAAAAGACTCAGGAAAAGGATATAACATCGTACAAGAAGGCTTTCGCCTGACTGTTGGTG<br>TAGACGCAGTTTGGGGTCCTAGTGCTCCGGATGCATGGGTACAACCGAAACAAATTTCGGAAAATACCTCGGCACAAGTTGCAAAT<br>AGTGCTGATGATGCTTTTATGGCTCTCGCAGAGAAGTTAAAAGCGTTATAAAGTACGAGGAGGTAAATATATATGGTTTTCACCCT<br>GGAAGACTTCGTTGGTGACTGGCGTCAGACCGCTGGTTACAACCTCGACCAGGTGCTAGAACAGGGTGGTGTTTCTTCTTTATTCC<br>AGAACCTGGGTGTAAGCGTTACCCCGATCCAGCGTATCGTTCTGTCTGGTGAAAACGGTCTGAAAATCGACATCCACGTTATCATC<br>CCGTACGAAGGTCTGTCTGGTGACCAGATGGGTCAGATCGAAAAAATCTTCAAAGTTGTTTACCCGGTTGACGACCACCACTTCAA<br>AGTTATCCTGCACTACGGTACCCTGGTTATCGACGGTGTTACCCCGAACATGATCGACTACTTCGGTCGTCCGTACGAAGGTATCG<br>CTGTTTTCGACGGTAAAAAAATCACCGTTACCGGTACCCTGTGGAACGGTAACAAAATCATCGACGAACGTCTGATCAACCCGGAC<br>GGTTCTCTGCTGTTCCGTGTTACCATCAACGGTGTTACCGGTTGGCGTCTGTGCGAACGTATCCTGGCTTAAATAATTATGTAACT<br>TAACACAGGACATCAAAAATGCTTAAAGAACTTTTAATGACAGAAGCTAAAACTATCGATGCTTCCGTCGCTCTTGACAGTATTTT<br>CGAATCAGTTCAGCTTTCTCCGGAAGCTAAAGCTAACTTCAGCACTGTATTCGAAGCAACAGTCAAAAAACACGCCGTAGCTCTGG<br>CAGAATCTCACATTGAAAAGATTGCTGAGAAAGCAGAAGAAAAAGTAGAAGAAGAAAAAGAAAAAGCTAAAGAAGAAGCTCGCGAA<br>GAGCTGAAAGAAGCTGCTTCTAAATACTTTGAC | Editing template for site-directed *nluc* insertion downstream of the prohead assembly protein gene of phage K1/K4 |
